# Spatial Single Cell Lipid-Transcriptomic Coupling Reveals Metabolic Niches in Glioblastoma

**DOI:** 10.64898/2026.07.13.738204

**Authors:** Tim F.E. Hendriks, Gert B. Eijkel, Martinus P.G. Broen, Ann Hoeben, Steven De Vleeschouwer, Ron M.A. Heeren, Eva Cuypers

## Abstract

Glioblastoma is characterized by spatial heterogeneity, with tumor-core and invasive-edge regions differing in cellular composition, transcriptional state, and metabolic context. Spatial transcriptomics has improved understanding of glioblastoma tissue organization. However, cellular transcriptional programs and metabolic interpretation remain poorly resolved. Here, single-cell matrix-assisted laser desorption ionization mass spectrometry imaging (MALDI-MSI) with spatial transcriptomics were integrated on the same tissue sections to map lipid and transcriptomic organization across matched tumor-core and invasive-region samples from 10 glioblastoma patients. Common region-dependent cellular organization along the core - versus invasive regions were identified after accounting for patient specific signatures. Tumor cores were enriched with astrocyte-like malignant and immune cell types, whereas invasive regions showed increased contribution from oligodendrocyte (progenitor cells). Despite these compositional differences, tumor and invasive cells had a shared transcriptional state space, indicating that regional identity is shaped by altered spatial organization of shared cell states. Transcriptional programs associated with proliferation, hypoxia, extracellular matrix remodeling, tumor associated macrophages and microglia (TAMs) phagocytic activity, T-cell infiltration, and lipid synthesis were spatially structured and differed between tumor and invasive compartments. MALDI-MSI revealed broad lipidomic remodeling across these regions. Tumor regions were enriched in membrane and storage lipid classes, whereas invasive regions showed relative enrichment of lipid species associated with membrane turnover. Integrating lipid and transcriptomic layers revealed spatial lipid-gene program coupling, with tumor cores showing stronger and more coherent coupling than invasive regions. TAM phagocytic activity co-localized with cholesteryl ester abundance, while lipid synthesis coupled to phosphatidylcholine-rich astrocyte-like tumor niches. Exploratory analysis indicated that MGMT promoter methylation may be associated with this altered lipid-transcriptional coupling, particularly in TAM-associated lipid-handling programs. Together, these findings imply that spatial coordination between lipid states and transcriptional programs is a key feature of glioblastoma metabolic heterogeneity.

## Introduction

Glioblastoma (GBM) is the most common and lethal primary malignant brain tumor in adults, marked by inter-and intra-tumoral heterogeneity covering malignant states and a complex immune ecosystem ^1, 2^. Large single-cell RNA sequencing (scRNA-seq) and molecular spatial studies continue to refine this complex disease ^3–5^. These studies also underscore that the high diversity of malignant and non-malignant cell types, and their changing cellular programs across GBM, remains a major obstacle to therapy ^3, 6–9^. Recent atlases highlight broad cellular compositions and dynamic state switching within IDH-wild-type GBM, emphasizing why regionally precise, cell-resolved measurements are needed ^10–12^. However, most spatial atlases still primarily describe transcriptional organization, whereas the metabolic implementation of these cell states within intact tumor niches remains less well resolved.

A clinically relevant subject is the transition from tumor core to the invasive edge, where tumor cells mingle with intact brain and adopt phenotypes linked to recurrence and treatment failure ^13, 14^. Spatial transcriptomic consistently detects cellular programs such as; hypoxia in tumor cores and ECM remodeling, lipid synthesis and more distinct immune cues toward the margin ^13, 15, 16^. The invasive margin is therefore not simply a diluted extension of the tumor core, but a mixed tissue state in which malignant cells interact with neural, glial, vascular, and immune compartments which continuously modify the tumor environment. However, most datasets lack single cell lipid readouts or rely on serial sections that complicate true one-to-one cell comparisons. Mapping the core to edge axis at a single-cell level with defined cell types, and doing so in a single section, is therefore essential to connect niche context with the molecular programs that could drive invasion and evasion.

Lipid metabolism is key to GBM biology. GBM rewires cholesterol metabolism, cholesteryl-ester storage, fatty-acid synthesis and desaturation, membrane composition, signaling programs, and therapy resistance ^17–21^. These lipid pathways are directly relevant to the spatial programs that define GBM: proliferating cells require membrane phospholipids, hypoxic cells remodel lipid storage and macrophage-lineage cells can process cholesterol-rich debris, myelin-derived lipids, and tumor-derived membrane material. Despite this, lipid remodeling has rarely been coupled to specific cell types in situ along the core-edge axis at single-cell resolution^22^.

Matrix assisted laser desorption ionization mass-spectrometry imaging (MALDI-MSI) is fitted to measure lipid remodeling and filling this knowledge gap. MALDI-MSI resolves spatial lipid heterogeneity directly in tissue and, at high spatial resolutions (5 µm), approaches the single cell resolution ^23, 24^. Practical integration with high-plex RNA methods has been limited by co-registration challenges, especially when modalities are acquired on different serial tissue sections ^25^. Emerging research emphasizes single-section, co-registered multimodal workflows to combine lipids to the correct cells and neighborhoods without intra-section variability and geometric uncertainty ^26^. Such approaches make it possible to study the local coupling of transcriptional programs and lipid states, rather than treating transcriptomics and lipidomics as parallel but spatially disconnected molecular layers.

In this study, thin tissue sections from resected material of ten glioblastoma patients were analyzed, sampling both tumor core and invasive edge zone separately. For each region, MALDI-MSI and Xenium spatial transcriptomics were performed on the same section to enable single-cell multimodal integration. Here, per-cell lipid features were coupled with gene programs and spatial context. Using a foundational spatial biology software tool, EscDat, for co-registration and integration, cell typing in matched core and invasive samples from human GBM was performed ^26^. This allowed the quantification of cell-type composition differences between regions, map within-cell-type transcriptomic and lipidomic shifts across the core-edge axis and link gene programs to lipid classes at single-cell resolution. Lipid-gene coupling differs according to O6-methylguanine-DNA methyltransferase (MGMT) promoter methylation status in the tumor was further explored. A clinically relevant biomarker of temozolomide (TMZ) response, to assess whether clinical subgroups may also differ in spatial metabolic organization. The focus on lipid changes to specific cellular identities and neighborhoods in the same section, delivered a mechanistic, spatially resolved view of GBM regional biology with translational clinical relevance.

## Materials and Methods

### Sample preparation – MALDI-MSI

Ethical approval for the study was obtained from the UZ/KU Leuven Ethical Review Committee of University Hospitals Leuven (Gasthuisberg, Leuven, Belgium) under reference number S60290, and the project was registered with the UZ Leuven Tissue Bank. Informed consent was obtained from all patients prior to inclusion, permitting the use of resected GBM tissue for research purposes, including assessment of clinically relevant genetic and epigenetic markers. All experiments involving human material were conducted in accordance with institutional and ethical guidelines. GBM tissue samples (n = 10) were collected immediately following surgical resection at the Department of Neurosurgery, KU Leuven. Post-excision, specimens were snap-frozen and stored at-80°C until processing. For each patient, a tumor core and invasive region sample were collected separately. The invasive-region sample from patient 8 was lost during sample processing and was therefore excluded from further analysis. For cryo-embedding, tissues were embedded in a 10% gelatin – 2% carboxymethylcellulose (Sigma-Aldrich) matrix. Serial cryosections of 10 µm thickness were prepared using a Leica CM1850 cryostat (Leica Biosystems) in accordance with 10x Genomics Protocol CG000579. Sections were stored at-80°C until analysis. Prior to MALDI-MSI, sections were equilibrated to room temperature in closed boxes containing hygroscopic desiccant beads, followed by vacuum desiccation for 20 minutes. Matrix application was performed by sublimation of 50 mg of 2,5-dihydroxybenzoic acid (2,5-DHB) (Sigma-Aldrich) at 160°C for 160 seconds using an HTX Sublimator (HTX Technologies).

### MALDI-MSI

MALDI-MSI was performed on a timsTOF fleX instrument equipped with a microGRID stage (Bruker Daltonics GmbH, Germany). Spectra were acquired in positive ion mode across an *m/z* range of 250-1200, with a pixel size of 5 x 5 µm and a laser spot diameter of approximately 4 µm. The Nd:YAG 355 nm SmartBeam 3D laser (Ekspla, Lithuania) was operated at a frequency of 1 kHz, and 25 laser shots were accumulated per pixel. External time-of-flight (TOF) calibration was carried out prior to imaging using red phosphorus. Instrument parameters were optimized as follows: funnel 1 RF, 200 Vpp; funnel 2 RF, 350 Vpp; and multipole RF, 450 Vpp. The focus pre-TOF transfer time was set to 100 µs. The quadrupole ion energy was maintained at 5.0 eV with a low-mass cutoff of *m/z* 200. The collision cell energy was set to 10.0 eV, and the collision RF amplitude to 1500 Vpp.

### Spatial transcriptomics

Following MALDI-MSI acquisition, residual 2,5-DHB matrix was removed by washing the slides twice for 1 minute each in 100% ethanol (BioSolve). The slides were subsequently dried under a steady stream of nitrogen gas. The Xenium slides were then fixed and permeabilized following the 10x Genomics Protocol CG000581. As an additional step, slides were fully submerged in phosphate-buffered saline (PBS) at 37 °C for 2 minutes. Any remaining gelatin covering the Xenium slide fiducial markers was gently removed using laboratory tissues (KimTech wipes, Sigma-Aldrich). Probe hybridization, ligation, and amplification were performed using the pre-designed Human Brain reagent kit according to 10x Genomics Protocol CG000749. All Xenium reagents were purchased and handled as described in Protocol CG000601. The processed slides were subsequently loaded into the Xenium Analyzer in accordance with Protocol CG000584. Following imaging and transcript capture, raw FASTQ files were processed using Xenium Ranger (version 2.0.0.10) with the Xenium multimodal cell segmentation workflow. The resulting data were visualized in Xenium Explorer (version 3.2.0) and further processed using the Seurat R package (version 5.3.0).

### LC-MS/MS acquisition and lipid identification

For lipid identification, approximately 5 mg of tissue from the tumor core and invasive regions of human GBM samples was manually resected and used for lipid extraction. Tissue was homogenized in 100 µL methanol (BioSolve), followed by addition of 400 µL methyl-tert-butyl ether (MTBE; Sigma-Aldrich). Samples were vortexed for 1 min and incubated in a thermoshaker for 1 h at 20°C and 950 rpm. Phase separation was induced by adding 100 µL water (BioSolve), followed by vortexing for 1 min and centrifugation at 1000 x g for 10 min using an Eppendorf 5353 centrifuge. The lipid-containing upper organic phase was transferred to a new tube. The remaining lower phase was re-extracted by adding 300 µL MTBE, 40 µL methanol, and 30 µL water, followed by vortexing and centrifugation under the same conditions. The second upper organic phase was combined with the first extract, dried in a vacuum centrifuge (Heto Lab), and reconstituted in 50 µL 1:1 isopropanol:acetonitrile (BioSolve). LC-MS/MS analysis was performed using a UHPLC system coupled to an Orbitrap Elite mass spectrometer (Thermo Fisher Scientific). Lipid separation was performed using a Hypersil GOLD column (100 x 2.1 mm). Data-dependent acquisition (DDA) LC-MS/MS measurements were performed in positive ionization mode ^27^. MS1 spectra were acquired over an *m/z* range of 200-1450 at a mass resolution of 60000 and an injection time of 65 ms. MS2 spectra were acquired in the ion trap using collision-induced dissociation (CID), an isolation window of 1.7 Da, and a mass resolution of 30000. LC-MS/MS lipid species were assigned using MS1 and MS2 spectra in Lipostar 2 version 2.1.7 using the LIPID MAPS database. MALDI-MSI lipid annotations were assigned by matching MS1 *m/z* values detected by MALDI-MSI to lipid identifications supported by MS1 and MS2 information from LC-MS/MS, using a mass tolerance of 15 ppm.

### Integration of MALDI-MSI and spatial transcriptomics

Single cells detected via the Xenium multimodal cell segmentation were converted to GeoJSON containing the cell identifier, per-gene transcript counts and spatial coordinates using a custom Python script. MALDI-MSI and Xenium datasets were then co-registered and integrated using EscDat (Version P1, 04-06-2025; MATLAB R2022b)^26^. A total ion current (TIC) image was generated and matched it to the fluorescence microscope image produced during Xenium’s multimodal cell-segmentation staining. MALDI laser ablation spots are visible in the fluorescence channel and were used, together with tissue fiducials placed prior to MALDI-MSI for coregistration. A piecewise-linear transformation from these landmarks yielded a transformation matrix linking the MSI and Xenium coordinate frames. MALDI-MSI peaks were preserved with a TIC-normalized intensity ≥ 5, and pixel intensities were weighted by the fractional overlap with each cell’s segmentation mask to mitigate partial-volume effects ^28^. EscDat was used for both the co-registration and the per-cell data integration steps.

The resulting integrated output was exported as a.csv file containing, for each cell, the cell identifier, spatial coordinates, transcript counts for each measured gene feature, and MALDI-MSI intensity values for each retained mass feature.

### Data analysis

Integrated per-cell Xenium-MALDI-MSI data were imported into Seurat for downstream analysis ^29–31^. Cell-type annotation was performed prior to gene program scoring using Xenium transcript counts. Cells were annotated by anatomical region (“Tumor” or “Invasive”) and patient identity. Cells were clustered based on their gene-expression profiles and annotated by the expression of canonical marker genes (*supplementary table S1*) ^32–44^. Marker-based annotations were then inspected both in RNA-based low-dimensional embeddings, such as UMAP, and in the original spatial coordinates of the tissue sections to ensure that annotated cell types showed expected transcriptional clustering and biologically plausible tissue distributions. To quantify region-associated differences in transcriptional program activity between tumor-core and invasive-region samples, a patient-level spatial index based on spatial transcriptomics-derived gene program scores was derived. Gene program scores were calculated at single-cell resolution from normalized Xenium transcript counts using predefined marker-gene sets for proliferation, hypoxia-associated adaptation, ECM remodeling, TAM phagocytic activity, T-cell infiltration and lipid synthesis. For each cell and each gene program, the average normalized expression of the genes belonging to that program was calculated relative to a background set of control genes with comparable expression levels following the Seurat module-scoring approach. The resulting score therefore represents the relative enrichment of a transcriptional program within an individual cell rather than the absolute expression of a single marker gene. Program scores were stored as metadata in the Seurat object and subsequently summarized by patient, anatomical region and cell type for tumor-core versus invasive-region comparisons. For each patient and each gene program, a tumor-core minus invasive-region difference score was calculated by subtracting the mean program score in the invasive region from the mean program score in the tumor core per patient. These patient-specific delta values therefore captured the direction and magnitude of regional program enrichment, with positive values indicating higher program activity in the tumor core. To combine multiple programs into a single patient-level measure, delta values were standardized across patients using z-score normalization, ensuring that each program contributed equally regardless of its absolute score range. The spatial core index was then defined as the mean of these standardized delta values. Higher spatial core index values indicate stronger divergence for tumor core and invasive regions, reflecting a more pronounced core-like transcriptional phenotype. Lower values indicate reduced spatial differentiation or a more invasive edge like state. Patients lacking measurements for either tumor or invasive regions were excluded from analyses requiring delta computation but retained in descriptive summaries.

For lipid-based dimensionality reduction, TIC-normalized MALDI-MSI intensities were used to generate a principal component representation of the per-cell lipid profiles. Harmony normalization was then applied to reduce inter-patient batch effects and/or patient specific signatures while preserving region-associated variation. The corrected lipid representation was visualized using UMAP, producing a low-dimensional embedding in which cells with similar lipid profiles are positioned near each other. Differential lipid abundance between tumor-core and invasive-region samples was assessed using integrated per-cell MALDI-MSI data. Lipid intensities were first compared at the regional level by summarizing all cells within each patient and anatomical region and subsequently at the cell-type level by aggregating intensities within each patient, region and annotated cell type. Analyses were restricted to lipid features with LC-MS/MS-supported annotations matched to MALDI-MSI m/z values within a 15 ppm mass tolerance. For each annotated lipid feature, MALDI-MSI intensities were assigned to individual cells based on the fractional overlap between MSI pixels and Xenium-derived cell segmentation masks. Lipid intensities were then summarized by anatomical region, patient, cell type and lipid class to compare tumor-core and invasive-region lipid profiles. Regional lipid differences were evaluated at both the lipid-class and individual lipid-species levels and cell-type-resolved tumor-invasive differences were calculated. Annotated lipid species were prioritized for visualization to facilitate biological interpretation. Given the limited cohort size, analyses were interpreted based on effect size, consistency across patients and cell types and spatial concordance rather than statistical significance alone.

## Results

### Patient cohort

Ten adults with IDH-wildtype glioblastoma, each contributing a tumor-core section and a patient-matched invasive-zone section, were profiled. In total, 19 sections were retained for analysis after exclusion of one invasive-zone section. The cohort included seven males and three females, with ages ranging from 32 to 73 years (median 64.5 years, mean 60.5 years). MGMT promoter status (Infinium MethylationEPIC BeadChip, Illumina) separated the cohort into five MGMT-methylated and five MGMT-unmethylated cases, enabling an exploratory comparison of lipid and transcriptional organization by clinically relevant molecular status. Given the limited cohort size, MGMT-associated analyses were interpreted as hypothesis-generating rather than definitive subgroup comparisons. For every retained section, MALDI-MSI and Xenium spatial transcriptomics were acquired on the same tissue section, enabling spatially aligned lipid and transcriptomic measurements at single-cell resolution. This design allowed direct assignment of cell identity, anatomical region, transcriptional program activity, and MALDI-MSI derived lipid intensity within the same tissue context. Across the retained sections, 198137 spatially resolved cells were analyzed, including 122707 cells from tumor-core regions and 75430 cells from invasive regions. The paired tumor-core and invasive-zone sampling further enabled patient-controlled assessment of spatial differences along the core-invasive axis. The full patient-level spatial maps (*supplementary figure 1*) show inter-patient heterogeneity in tissue architecture and cell-type density, while preserving regional patterns. Patient-level clinical metadata retained regions and cell numbers are summarized in *supplementary table S2*.

### Cell typing and composition core vs invasive

A spatially resolved cellular framework was established for subsequent multimodal lipid-transcript integration and defined how cell-type composition shifts along the glioblastoma core-to-edge axis between patients. Since lipid availability is strongly shaped by local cellular neighborhoods, precise quantification of regional cellular composition is essential to interpret spatial lipid remodeling mechanistically. Leveraging single-section co-registered Xenium profiling across matched tumor core and invasive regions from ten patients, a per-cell map without inter-section alignment bias was constructed. Cell-type annotation was supported by canonical marker-gene expression (*supplementary table S1 and supplementary figure 2)*, including AQP4, FGFR3, and GJA1 for astrocyte-like cells; MOBP, MOG, and OPALIN for oligodendrocytes; OLIG2, PDGFRA, and SOX10 for oligodendrocyte progenitor cells (OPCs); CD163, CD68, CX3CR1, P2RY12, and TREM2 for TAMs; CD4 for T-cell associated cells; PECAM1, FLT1, and NRP1 for endothelial cells; CRYM, SLC17A6 and SLC17A7 and GAD1, SST and WIF1 for excitatory and inhibitory neurons; and PAX6/SOX2 and MGST1/NCSTN/TGFB2 for glioblastoma stem cells (GSC) and mesenchymal (MES)-like populations respectively ^32–44^. Patient-level cell counts for each retained tumor-core and invasive-region sections are provided in *supplementary table S3* and were used as the basis for composition and paired tumor - invasive cell-type abundance analyses. Tumor-core sections frequently showed dense astrocyte-like malignant regions together with immune-lineage enrichment, primarily involving TAMs and, in some patients, T-cell-containing niches. The relative contribution of MES-like, GSC-like, OPC-like, and immune populations varied between patients. Invasive-region sections showed a greater relative contribution of oligodendrocyte and neural-lineage populations, consistent with infiltration into preserved brain parenchyma rather than a uniform region-specific cell identity. This spatial overview confirms that the tumor core-invasive comparison captures both the shared cell identities and region-dependent changes in cellular organization. Representative spatial maps (Figure 1A) illustrate that both compartments contain highly intermixed malignant and stromal populations, a pattern expected in GBM given its diffuse growth and cellular heterogeneity. The single-section integrated workflow enabled direct visualization of these compositional gradients within intact tissue architecture, highlighting that the invasive niche is not simply diluted tumor mass but a structurally distinct microenvironment in which malignant cells coexist with preserved healthy elements. Quantitative assessment of cell-type fractions per patient and region (Figure 1B) revealed consistent regional biases despite inter-patient heterogeneity. Astrocyte-like cells were a dominant fraction within tumor cores across most cases, whereas their proportional contribution decreased in invasive regions. In contrast, oligodendrocytes reproducibly increased at the invasive edge. MES states and glioma stem-like cells (GSCs) were variably distributed but tended to be relatively enriched in tumor cores. Immune and vascular compartments displayed patient-dependent patterns. Paired analyses of cell-type abundance differences between invasive-region and tumor-core samples confirmed the regional trends while also revealing substantial inter-patient variability (Figure 1C). Positive values indicate invasive enrichment, whereas negative values indicate tumor-core enrichment. Oligodendrocytes showed a significant positive difference, indicating invasive enrichment, whereas astrocyte-like cells showed a robust negative paired difference consistent with tumor-core predominance. MES-like cells and GSCs showed more modest and heterogeneous shifts. Stem-like programs concentrate within core niches although they are not strictly compartmentalized. In contrast, both CD4 positive T-cells and TAMs indicate relative enrichment within tumor cores rather than at the invasive margin. This implies that immune cell accumulation is more pronounced in the core microenvironment. The patient-level heatmaps (Figure 1D) further highlighted the reproducibility of astrocyte depletion and oligodendrocyte enrichment at the invasive edge across the cohort, while immune and endothelial changes were more heterogeneous.

**Figure 1.**
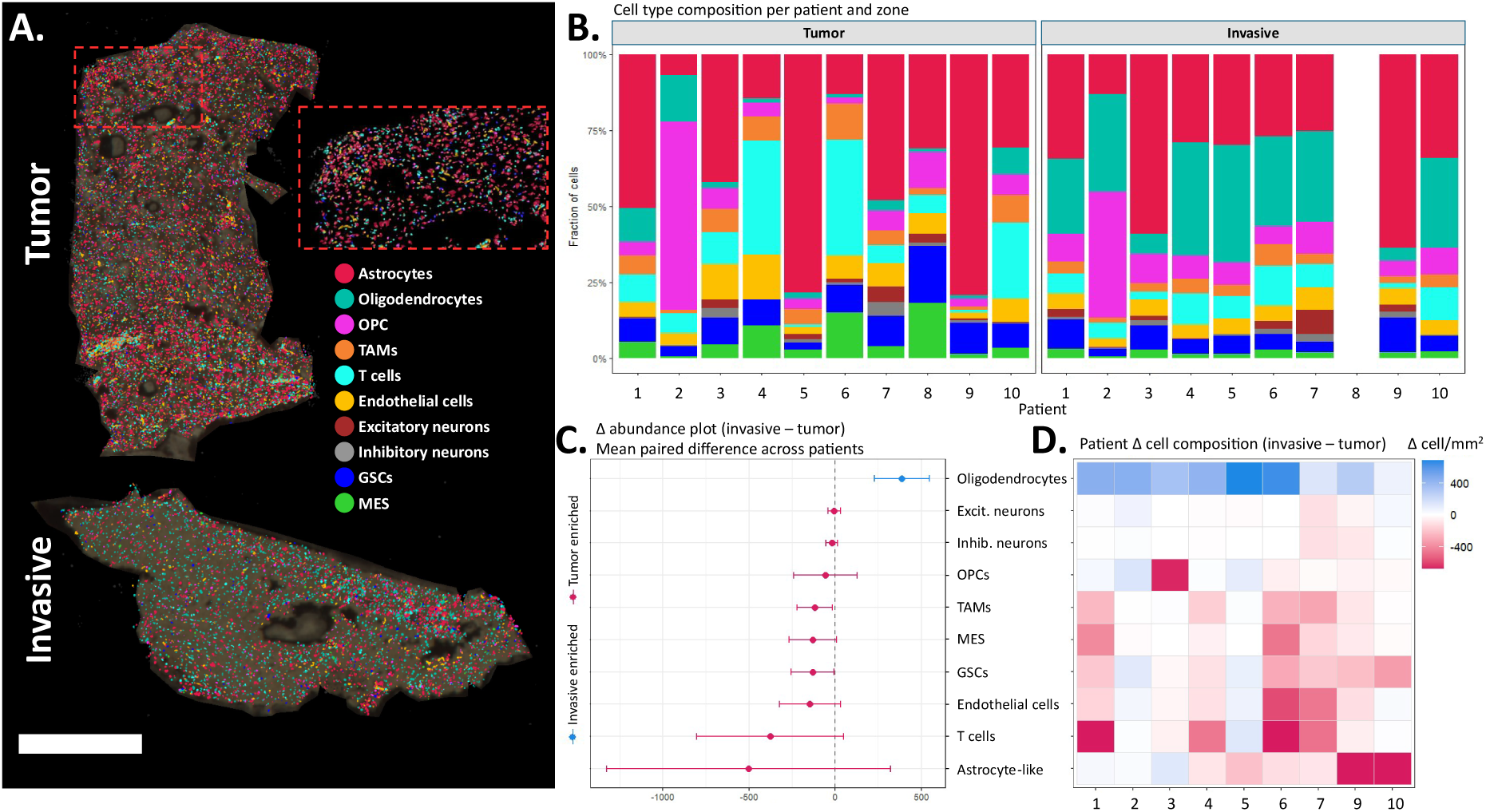
**Spatially resolved cell-type composition differs between glioblastoma tumor core and invasive regions across patients**. **(A)** Representative Xenium single-cell spatial transcriptomics maps from one patient (patient 10) showing the tumor core (top) and invasive region (bottom). Individual cells are colored by cell-type annotation; dashed box indicates a zoom-in region highlighting local cellular intermixing. Scale bar shows 1 mm. **(B)** Cell-type composition per patient and region, shown as fractions of density (cells per mm²). Patient 8 had no invasive sample **(C)** Forest plot summarizing paired differences in cell-type abundance between invasive and tumor regions across patients (Δ = invasive - tumor). Points indicate the mean paired difference, and horizontal bars represent 95% confidence intervals. Positive values show invasive-enriched cell types, while negative values show tumor-enriched cell types. **(D)** Patient-level heatmap of cell-type abundance differences (Δ = invasive - tumor; cells per mm²). Columns are patient numbers.

A joint UMAP embedding of all RNA-profiled cells (*supplementary figure S3*) further shows that tumor and invasive cells occupy the same transcriptional state, indicating that invasive cells do not form distinct region-specific clusters. Instead, the primary regional difference lies in the relative occupancy of shared cell-type populations. Tumor-core samples were generally characterized by increased astrocyte-like malignant populations and patient-dependent enrichment of immune and malignant-state compartments, including TAMs, T-cells, GSCs, and MES-like cells. Invasive-region samples showed a more consistent increase in oligodendrocyte populations. These differences provide the need for subsequent analyses of region-associated transcriptional programs within cell types, including proliferation and cell-cycle activity, hypoxia-associated metabolic adaptation, extracellular matrix and lipid remodeling linked to invasion.

### Transcriptomic reprogramming within cell types across the tumor core - edge axis

The prior analysis revealed consistent differences in cellular composition between tumor-core and invasive-region samples. However, compositional shifts alone do not explain how shared cell populations adapt to distinct microenvironmental contexts. Cells of the same lineage can adopt region-dependent states in response to local environments such as oxygen availability, extracellular matrix organization, immune activity and metabolic demand. We therefore next investigated whether the core and invasive axis is accompanied by transcriptional reprogramming within our annotated cell types.

To quantify these region dependent states, six transcriptional programs were defined representing proliferation and cell-cycle activity, hypoxia-associated metabolic adaptation, extracellular matrix (ECM) remodeling, microglia-macrophage phagocytic activity, T-cell infiltration and lipid synthesis (Figure 2A). The genes used to define each program are also provided in *supplementary table S4*. Expression of these program defining genes across annotated cell types confirmed expected lineage associations. Proliferation-associated genes were enriched in malignant and stem cell-like compartments, hypoxia and ECM-associated genes were distributed across malignant and stromal populations. The TAM phagocytic program showed its strongest signal in TAMs, consistent with a macrophage-lineage-associated phagocytic/lipid-handling state. Genes within the T-cell infiltration program were most prominent in the annotated T-cell population, but several genes in this module also showed signal in TAM-rich regions. This module was interpreted as reflecting broader immune-niche activity, including lymphoid presence and myeloid-associated inflammatory signaling, rather than a pure T-cell lineage or activation-state marker. Lipid synthesis genes were broadly expressed across multiple cellular populations. Together, these patterns show that the selected gene sets do not solely reflect isolated marker expression, but capture coordinated transcriptional programs that map onto biologically relevant cell states. This provides a functional framework for comparing how malignant, immune, stromal and neural populations adapt across the tumor-core and invasive-region microenvironments.

**Figure 2.**
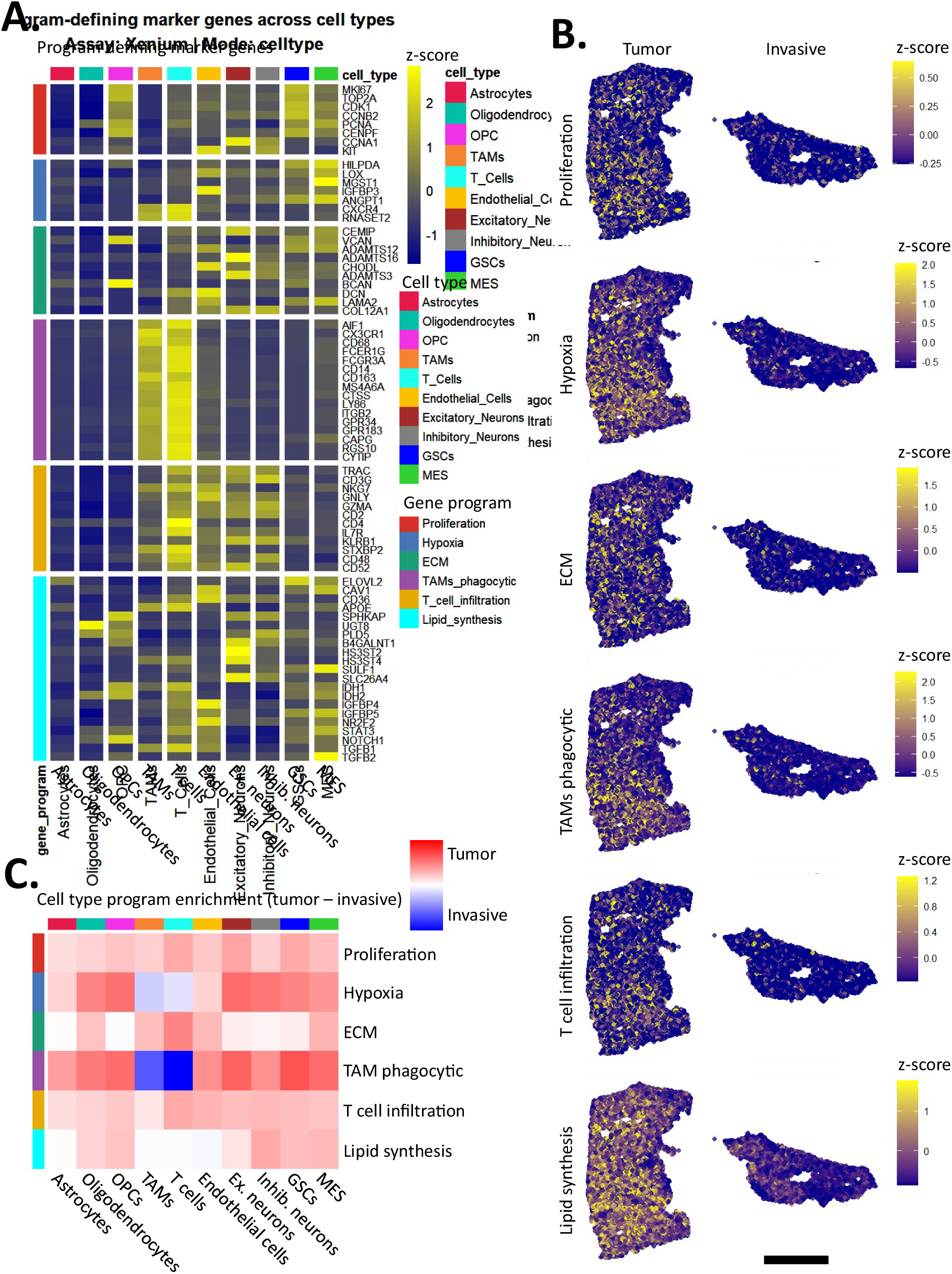
Spatially organized transcriptional programs across the glioblastoma tumor core-invasive axis. **(A)** Heatmap showing scaled expression of program-defining marker genes across annotated cell types. Rows represent genes grouped into six transcriptional programs: proliferation, hypoxia-associated adaptation, ECM remodeling, TAM phagocytic activity, T-cell infiltration, and lipid synthesis. Columns represent annotated cell types. Yellow indicates higher scaled expression, and blue indicates lower scaled expression. **(B)** Spatial maps of transcriptional program scores in representative matched tumor-core and invasive-region sections (patient 10). Each point represents a single cell colored by z-scored program activity. Color scales are program-specific, with higher values indicating increased score activity. Scale bar shows 1 mm. **(C)** Heatmap summarizing cell-type-specific regional differences in transcriptional program activity. Values represent tumor-core minus invasive-region program scores for each cell type and program. Red indicates higher activity in tumor-core samples, blue indicates higher activity in invasive-region samples, and white indicates little regional difference.

Spatial visualization of program scores within a representative patient (patient 10) showed that these transcriptional states were not uniformly distributed across tissue but instead formed regionally structured patterns (Figure 2B) that aligned with the cellular architecture defined in Figure 1. In the representative tumor-core section, proliferation, hypoxia, ECM remodeling, TAM phagocytic activity, T-cell infiltration and lipid synthesis showed spatially heterogeneous activity, with localized areas of increased program score rather than homogeneous activation across the tissue. Proliferation, hypoxia and lipid synthesis programs were broadly distributed within tumor-core tissue, whereas immune-associated programs were more focal, with TAM phagocytic and T-cell infiltration signals appearing in discrete niches. In the matched invasive-region section, these same programs were still detectable but showed less focal organization and generally lower local enrichment. Rather than showing one dominant invasive-activated program, the invasive region displayed more diffuse and cell-type-dependent program activity, consistent with its altered cellular composition and lower density of tumor-associated cell populations.

Next, regional program differences across cell types by comparing tumor-core and invasive-region program activity were summarized (Figure 2C). This analysis revealed that most programs showed cell-type-specific shifts rather than uniform region-wide changes. Proliferation, hypoxia, ECM remodeling, T-cell infiltration and lipid synthesis generally showed higher activity in tumor-core samples across several cell types, although the magnitude varied by cell type. In contrast, the TAM phagocytic program displayed a more cell-type dependent regional pattern, with the strongest signal in the T-cell and TAM compartments. This program should be interpreted as a macrophage-lineage signature related to phagocytosis and lipid/debris processing, rather than as evidence of an anti-tumoral TAM phenotype.

To determine whether these program-level differences were driven uniformly by all genes within each module, the tumor-core versus invasive-region differences at the individual gene level was also examined (*supplementary figure S4*). This analysis showed that the summarized program scores in Figure 2C reflect coordinated but not homogeneous gene-level behavior. This indicates that the GBM core-invasive axis is shaped not only by changes in cellular composition, but also by selective transcriptional reprogramming within shared cell types. Because several of these programs (including hypoxia-associated adaptation, ECM remodeling, TAM phagocytic activity, and lipid synthesis) are directly linked to metabolic stress, membrane remodeling and lipid handling, it was investigated whether these transcriptional states are accompanied by changes in lipid composition. To address this, transcriptomic program scores were integrated with spatial lipid measurements obtained by MALDI-MSI from the same tissue sections, enabling direct cell-resolved comparison of gene expression programs and lipid distributions.

### Lipidomic reprogramming within cell types

Having established that transcriptional programs exhibit spatial and cell-type specific organization across the tumor core-edge axis, it was asked whether these changes are accompanied by coordinated changes in lipid composition. Annotated lipid features were matched to LC-MS/MS-derived lipid identities and curated by lipid class and summarized in *supplementary table S5*. A low-dimensional UMAP embedding of the MALDI-MSI lipid profiles was generated after Harmony-based normalization to reduce inter-patient batch effects. In this lipid-based embedding, tumor-core and invasive-region cells separated clearly and indicate that MALDI-MSI lipid profiles alone capture regional differences between these compartments (Figure 3A). Prior to batch correction, lipid features were strongly segregated by patient identity, reflecting substantial inter-sample variability inherent to MALDI-MSI data (*supplementary figure S5*). Harmony based normalization effectively removed this patient-specific signal while preserving relevant variation, enabling direct comparison across samples ^45, 46^. The resulting segregation of tumor and invasive zones (Figure 3A) demonstrates that the core-edge transition is accompanied by metabolic differences, extending on transcriptional differences to the level of lipid composition.

**Figure 3.**
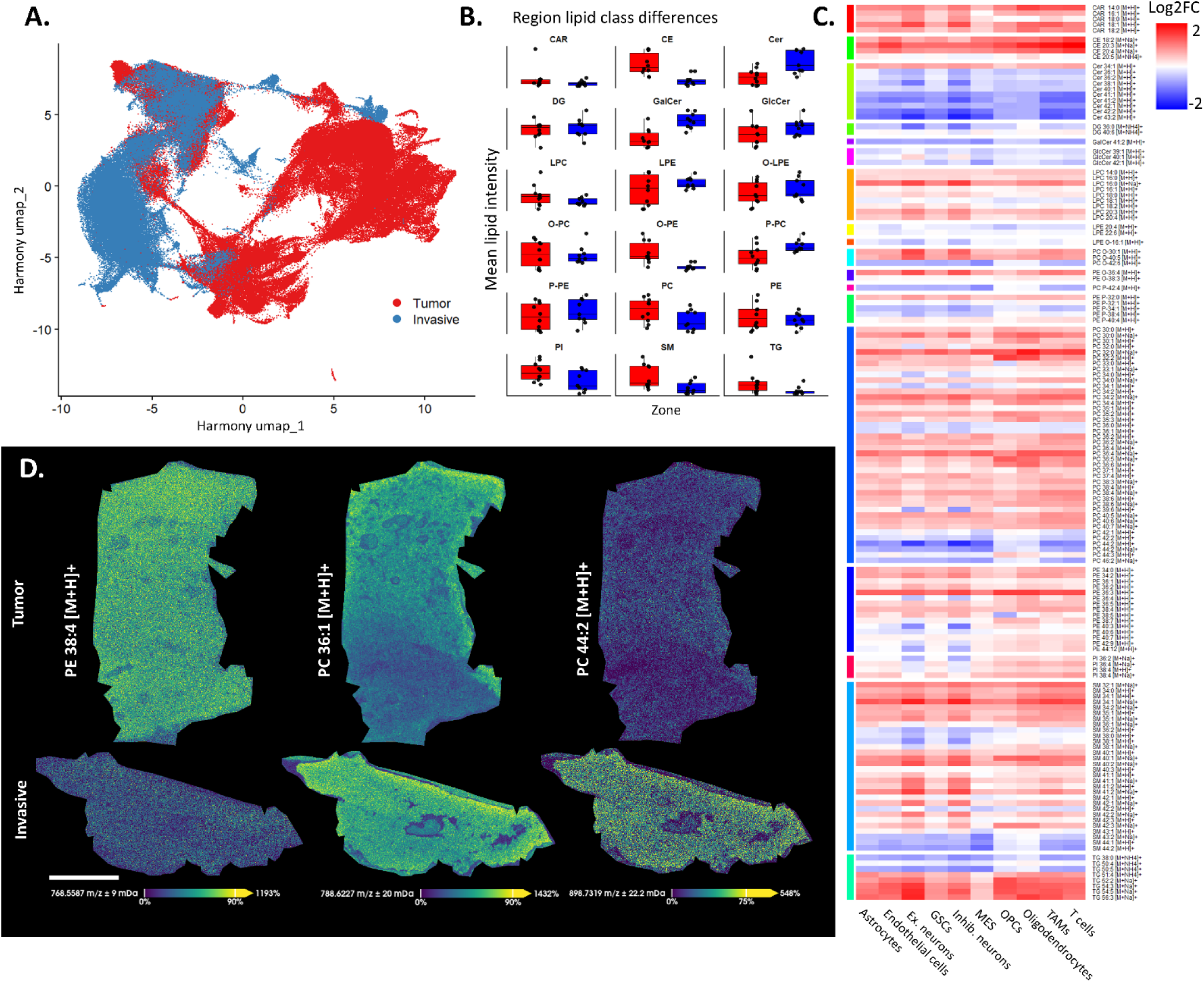
Spatial lipidomic reprogramming across the glioblastoma core-edge axis. **(A)** Unsupervised embedding of MALDI-MSI lipid features after Harmony batch correction separates tumor core and invasive regions. **(B)** Lipid class differences between tumor (red) and invasive (blue) regions. Tumor cores are enriched in membrane and storage lipids, including phosphatidylcholines (PC), phosphatidylethanolamines (PE), ether-linked phospholipids, sphingomyelins (SM), cholesterol esters (CE), and triglycerides (TG), whereas invasive regions show enrichment of lysophospholipids (LPC) and subsets of ceramides and glycosphingolipids. **(C)** Cell-type resolved differential lipid abundance (log2FC, tumor vs invasive) reveals structured, class-specific remodeling. PCs are consistently tumor-enriched across cell types, while PEs show heterogeneous, cell-state dependent patterns. Lysophospholipids are enriched in invasive regions, and sphingolipids display variable, lineage-specific regulation. **(D)** Spatial mapping of representative lipid species demonstrates that tumor-enriched phospholipids localize to tumor regions, with reduced abundance at the invasive margin. Representative section patient 10. Scale bar indicates 1 mm.

A class-level lipid abundances comparison between regions was performed (Figure 3B) to determine whether these differences reflect coordinated remodeling of specific lipid classes. Lipid classes, including phosphatidylcholines (PC), phosphatidylethanolamines (PE), their ether-linked counterparts (O-PC, O-PE), cholesterol esters (CE) and sphingomyelin (SM) showed consistent enrichment in tumor regions. In parallel, triglycerides (TG) associated with lipid storage and metabolic buffering, were also elevated in tumor cores. In contrast, lysophosphatidylcholines (LPC) and subsets of ceramiders (CE, GalCer, GlcCer) showed enrichment in invasive regions. These findings indicate that lipidomic differences between tumor and invasive compartments are not driven by isolated lipid species, but instead reflect coordinated class-level remodeling.

Analysis of differentially abundant lipid species revealed structured patterns that depended on both lipid class and cell type (Figure 3C). The PC class contained a large tumor-enriched lipid block, with many PC species showing positive log2FC across malignant, stromal, and immune populations. PC species showed a predominantly tumor-enriched pattern, despite some cell-type-specific variation. By contrast, PE and ether-linked PE species showed a more mixed regional pattern, with both tumor-core and invasive-region enrichment depending on lipid species and cell type. Indicating that PE remodeling is more cell type-and lineage-dependent than the broader PC pattern. Interestingly, PE 36:3 [M+H]+ was highly positive for tumor across all cell types. Invasive regions were characterized by enrichment of LPEs, which showed consistent negative log2FC across cell types, suggesting increased membrane remodeling at the tumor margin. Sphingolipids, sphingomyelins and glycosphingolipids, exhibited more variable, cell-type specific patterns, with notable differences in neuronal and oligodendrocyte populations. Ceramides also showed higher intensities in invasive regions. Triglycerides (TG) were predominantly enriched in tumor regions, particularly in TAMs, T-cells, OPCs and oligodendrocytes, consistent with increased lipid storage or metabolic buffering. Representative lipid species were visualized directly within tissue sections (Figure 3D) to validate these observations spatially. The tumor-enriched phospholipid, such as PE 38:4, displays a strong and spatially coherent signal within tumor regions, whereas the abundance was reduced in invasive areas. These spatial distributions validate the statistical findings from the lipid class-level and species-level analyses by demonstrating that lipids identified as tumor-enriched are also spatially concentrated within tumor regions of the tissue. Notably, individual lipid species exhibited distinct spatial gradients rather than uniform distributions, further supporting the presence of localized specific metabolic states within the tumor microenvironment.

### Spatial coupling of lipid states and transcriptional programs across the tumor core-edge axis

The data demonstrates that transcriptional programs and lipid composition both vary along the tumor core-edge axis. Therefore, whether these molecular layers are coordinated within the same spatial and cellular contexts were examined next. The region-associated transcriptional programs are connected to stress-associated cellular processes. However, transcriptional activity does not necessarily translate directly into downstream lipid abundancy changes, particularly in heterogeneous tissues where metabolic state and cell-cell interactions can differ across local niches. The per-cell transcriptomic program scores with co-registered MALDI-MSI lipid intensities were integrated to assess whether individual lipid species and lipid classes co-vary with specific gene programs across tumor and invasive regions. Here, lipid-gene coupling refers to the spatial covariation between transcriptional program activity and MALDI-MSI-derived lipid abundance across cells or local tissue regions. Positive coupling indicates that higher activity of a given gene program occurs together with higher abundance of a specific lipid class or lipid species. Weak coupling indicates little spatial relationship between the two molecular layers, whereas negative coupling indicates an inverse relationship. Lipid-gene coupling analysis revealed that transcriptional programs are associated with structured lipid signatures, meaning that specific gene programs reproducibly correlate with defined lipid classes or lipid species, rather than diffuse or random lipid variation. When correlations were calculated separately in tumor core and invasive regions, the tumor compartment showed a more organized pattern of coupling meaning that correlations were stronger, more coherent within lipid classes, and more consistently aligned with specific transcriptional programs. In contrast, the invasive region displayed more weak and heterogeneous associations (Figure 4A). This indicates that the relationship between transcriptional state and lipid composition is region dependent, with tumor cores probably exhibiting tighter coordination between gene programs and lipid metabolism. Notably, lipid classes involved in membrane composition, lipid storage, and lipid signaling showed distinct coupling patterns across programs, suggesting that different transcriptional states are linked to different lipidomic phenotypes. This is also seen in the correlation matrices including all annotated lipid species and all measured Xenium genes. This showed a broader class-structured lipid-gene organization in tumor-core samples, whereas invasive samples displayed less consistent correlation patterns (*supplementary figures S6 and S7*). The mean absolute gene-lipid coupling strength for each transcriptional program was calculated (Figure 4B) to summarize these relationships at the lipid-class level. Across multiple programs, coupling was generally stronger in tumor cores than in invasive regions, consistent with the more organized patterns observed in the correlation heatmaps. Proliferation, T-cell infiltration, and hypoxia showed particularly strong tumor-associated coupling with several lipid classes, whereas lipid synthesis and TAM phagocytic activity displayed a more variable pattern, with some lipid classes showing comparable or stronger coupling in invasive regions. Analysis of individual lipid species further demonstrated that transcriptional programs were associated with distinct lipid signatures (Figure 4C). Proliferation and lipid synthesis programs showed positive associations with several membrane-associated lipid species, including PC, PE, SM, and CE species. Hypoxia and ECM remodeling displayed more mixed lipid associations, with both positive and negative correlations across PC, PE, SM, and CE species. TAM phagocytic activity showed a prominent association with cholesteryl ester species, whereas T-cell infiltration was associated with sphingomyelin and phospholipid species. These findings indicate that lipid-gene coupling is program-specific rather than uniformly shared across all transcriptional states.

**Figure 4.**
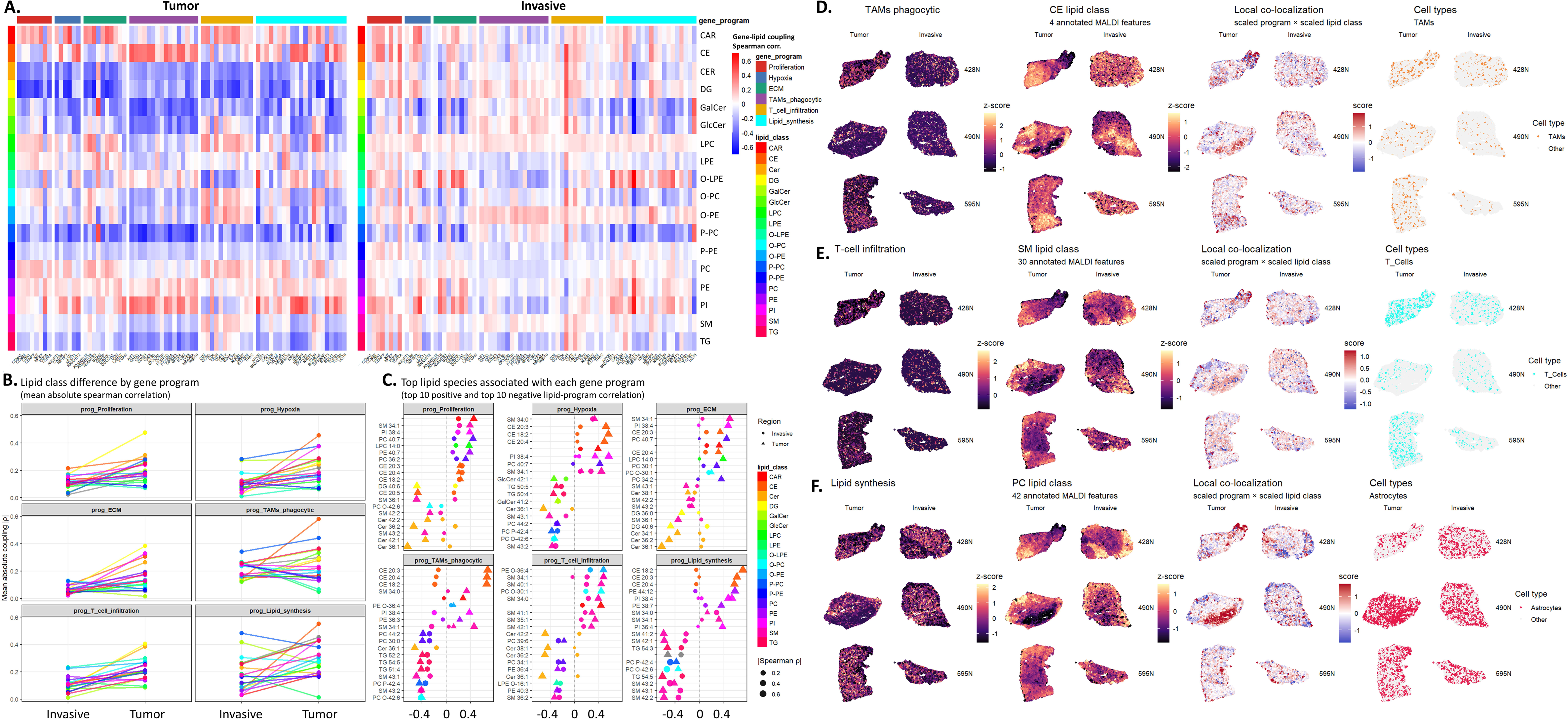
Spatial coupling between lipid states and transcriptional programs across the glioblastoma core-edge axis. **(A)** Lipid class resolved gene-lipid coupling analysis in tumor core and invasive regions. Heatmaps show Spearman correlations between MALDI-MSI lipid classes and genes grouped by transcriptional programs. **(B)** Summary of lipid class program coupling strength. Lines show mean absolute Spearman correlations for each lipid class in invasive and tumor regions across transcriptional programs. **(C)** Top lipid species associated with each transcriptional program. Dot plots show the strongest positive and negative lipid program correlations for each module, separated by region. Dot position indicates Spearman correlation coefficient, shape indicates tumor or invasive region, color denotes lipid class, and dot size represents correlation magnitude. Distinct transcriptional programs are associated with specific lipid species, supporting program-specific lipid remodeling across the core-edge axis. **(D)** Spatial validation (patient 6, 1 and 10 respectively) of TAM phagocytic program coupling with cholesteryl esters. Representative tumor and invasive sections from three patients show TAM phagocytic program scores, CE lipid class abundance, local co-localization between scaled TAM phagocytic score and scaled CE abundance, and spatial localization of TAMs. **(E)** Spatial validation of T-cell infiltration coupling with sphingomyelins. **(F)** Spatial validation of lipid synthesis coupling with phosphatidylcholines

Selected high-coupling pairs were mapped across matched tumor and invasive sections (Figure 4D-F) to determine whether the lipid-program associations were spatially organized within tissue. CE abundance showed focal co-localization with the TAM phagocytic program in regions containing TAMs, supporting a spatially restricted macrophage-associated lipid-handling niche (Figure 4D). Sphingomyelin abundance showed more heterogeneous but locally increased overlap with T-cell infiltration, indicating that immune-associated lipid coupling varies across patients and regions (Figure 4E). Finally, phosphatidylcholine abundance co-localized with lipid synthesis program activity in astrocyte-enriched areas, linking transcriptional lipid biosynthesis to local membrane phospholipid content (Figure 4F). Together, these spatial validations show that coupling is not driven solely by global regional differences but is organized into localized niches. Full-cohort spatial maps for all six transcriptional programs further supported this interpretation (*supplementary figures S8 - S13*). Across programs, local co-localization between program activity and the corresponding lipid class was spatially heterogeneous, with co-enriched regions observed across patients. These maps demonstrate that lipid-program coupling is not uniform across tissue, but varies by program, lipid class, and patient-specific architecture. These maps demonstrate that lipid-gene program coupling is not uniform across tissue, but varies by transcriptional program, lipid class and patient-specific tissue architecture. Tumor cores showed more spatially coherent associations between transcriptional program activity and MALDI-MSI-derived lipid abundance, whereas invasive regions showed weaker and more heterogeneous associations. This shows that lipid states and gene-program activity are more consistently coupled in tumor-core niches than in the invasive compartment.

### MGMT promoter methylation is associated with altered transcriptional-lipid coupling

Whether this multimodal organization is associated with clinically relevant molecular features of glioblastoma was investigated next. MGMT promoter methylation is one of the most important clinical biomarkers in glioblastoma and is routinely used to stratify response to temozolomide (TMZ)-based therapy. Therefore it was explored whether MGMT-methylated (MGMT^+^) and MGMT-unmethylated (MGMT^-^) tumors differ not only in their transcriptional programs or lipid abundance individually, but also in the extent to which lipid states are coupled to transcriptional programs. Because MGMT status is not expected to directly determine lipid metabolism, we treated this analysis as an exploratory clinical extension rather than as a primary consideration of metabolic state. First, lipid class abundance between MGMT-methylated and MGMT-unmethylated tumors across the cohort were compared (Figure 5A). Lipid abundance differences were present. CE showed the strongest increase in MGMT^+^ tumors, followed by diacylglycerols (DG). In contrast, several lipid classes, including PC, acylcarnitines (CAR), galactosylceramides (GalCer), LPE, LPC, PE and ceramides, were relatively higher in MGMT^-^ tumors. These data imply that MGMT status is not associated with a uniform increase or decrease in lipid content, but instead with selective remodeling of specific lipid classes, most prominently CE enrichment in MGMT^+^ tumors. Whether MGMT status was associated with tumor-invasive differences in transcriptional program activity was also assessed (Figure 5B). Most program differences showed modest separation between MGMT groups, but the largest MGMT associated shift was observed for the TAM phagocytic program, which was more strongly increased in MGMT^+^ tumor cores relative to invasive regions. Hypoxia and lipid synthesis also showed positive shifts in MGMT^+^ tumors, whereas proliferation and ECM remodeling were more variable and showed smaller effect sizes. These findings indicate that MGMT status does not broadly stratify all transcriptional programs but may be associated with enhanced immune and lipid associated programs in the tumor core.

**Figure 5.**
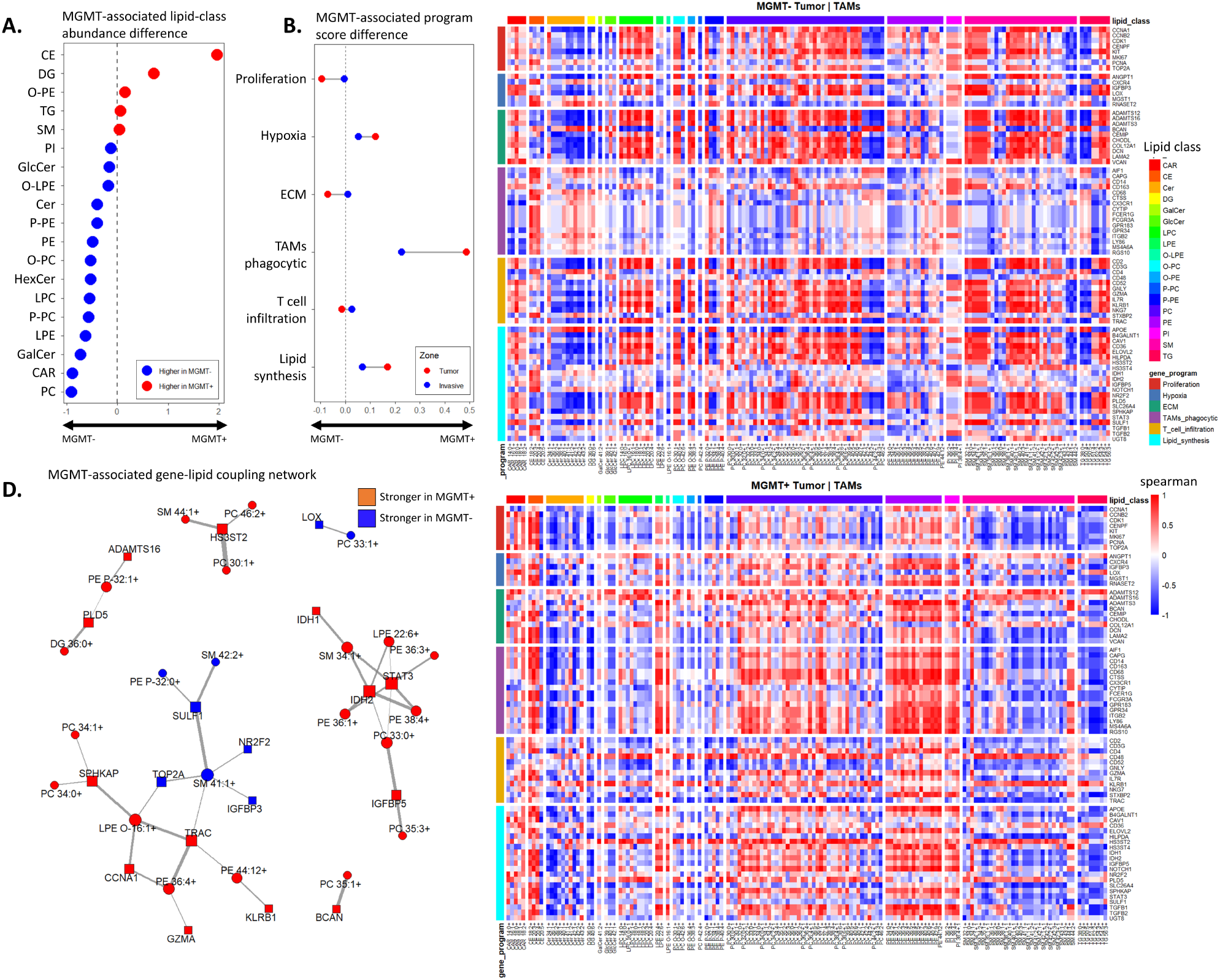
MGMT promoter methylation is associated with altered lipid-gene coupling. **(A)** MGMT-associated lipid-class abundance differences. Red indicates enrichment in MGMT+ tumors and blue indicates enrichment in MGMT-tumors. **(B)** MGMT-associated differences in GBM tumor-invasive transcriptional program scores. Red points represent tumor regions and blue points represent invasive regions. **(C)** Heatmaps showing MGMT-stratified gene-lipid coupling in TAMs. Program-associated genes and lipid species grouped by lipid class. Red indicates positive gene-lipid correlation, blue indicates negative correlation, and white indicates weak or no correlation. Spearman correlation. **(D)** Network of MGMT gene-lipid coupling differences. Red indicates associations stronger in MGMT+ tumors, whereas blue indicates associations stronger in MGMT-tumors. Spearman correlation.

Because lipid abundance and transcriptional program differences alone showed selective rather than global MGMT-associated effects, it was examined whether MGMT status altered the coupling between genes and lipids. Focusing on TAMs in tumor regions, gene-lipid correlation matrices revealed distinct coupling architectures between MGMT tumors (Figure 5C). In MGMT^-^ TAMs, lipid-gene correlations appeared strongly organized across multiple program-associated genes, with strong coupling involving sphingolipid, phospholipid, and TG classes. In contrast, MGMT^+^ TAMs showed a different correlation structure, including pronounced coupling between selected gene modules. These patterns could state that MGMT status is associated less with simple abundance differences and more with reorganization of how lipid species relate to transcriptional programs within macrophage-lineage cells. To further summarize this association, an MGMT-associated gene-lipid coupling network was constructed (Figure 5D). This network highlighted lipid-gene relationships that were stronger in either MGMT^+^ or MGMT^-^ tumors. MGMT^+^ tumors showed stronger coupling between multiple lipid species and genes related to immune signaling, extracellular matrix remodeling, and stress-associated programs, including relationships involving PE 36:3, PE 38:4, PC 33:0, LPE 22:6, SM 34:1, STAT3, IDH1, IDH2, and IGFBP5. In contrast, MGMT^-^ tumors showed stronger coupling involving selected sphingomyelin species with genes such as LOX, TOP2A, SULF1, NR2F2, and IGFBP3. This network representation supports the conclusion that MGMT status influences the organization of transcriptional-lipid relationships rather than simply shifting individual gene programs or lipid classes in isolation. This suggests that MGMT promotor methylation is associated with selective remodeling of lipid abundance and transcriptional program activity, with the strongest signal emerging at the level of lipid and gene coupling. In particular, MGMT^+^ tumors showed evidence of enhanced CE abundance and stronger tumor-associated TAM phagocytic program differences, accompanied by distinct lipid-gene coupling architecture in TAMs. Although these findings require validation in larger cohorts, they suggest that MGMT status may reflect not only treatment sensitivity but also differences in the spatial metabolic organization of the GBM microenvironment with main effect on TAM phagocytic programs.

## Discussion

In this study, we utilize a single section multimodal workflow that integrates MALDI-MSI lipid imaging with Xenium spatial transcriptomics to resolve the glioblastoma tumor core-edge transition at a single cell spatial resolution. By acquiring both modalities from the same tissue section, lipid features were directly combined to spatially annotated cell types and transcriptional programs, avoiding the uncertainty introduced by serial-section measurements. This design allows us to move beyond regional bulk comparisons and address a central question in GBM biology: How are cellular composition, transcriptional state and lipid metabolism coordinated within the spatial niches that define tumor growth and invasion ^24–26^?

Our data supports a model in which the tumor core and invasive region represent distinct but connected cellular ecosystems. Tumor cores were variably enriched for astrocyte-like malignant populations and immune-rich areas, including TAMs and T-cell containing regions, whereas invasive samples showed increased contribution from oligodendrocyte and OPCs, consistent with infiltration into preserved brain parenchyma and with prior work linking GBM invasion to neural and progenitor-like tumor states ^7^. This pattern is consistent with the concept that GBM invasion occurs through intermixing with preserved brain parenchyma, rather than through the emergence of a completely independent invasive cell state ^7, 10, 13, 16^. At the same time, invasive-region specimens inherently contain variable mixtures of infiltrating tumor and relatively preserved brain tissue. While this reflects the clinical reality of the invasive margin, it also increases heterogeneity and may contribute to variability in pseudobulk comparisons. In line with prior single-cell and spatial studies, tumor and invasive cells occupied shared transcriptional state space, indicating that regional identity is largely shaped by altered occupancy and spatial organization of shared cellular states ^3, 6, 47, 48^. The transcriptional program analysis further refines this view. Proliferation, hypoxia, ECM remodeling, TAM phagocytic activity, T-cell infiltration and lipid synthesis are spatially organized across tissue but are not uniformly restricted to single cell types or anatomical compartments. Instead, these programs showed activity across malignant, immune and neural cell populations. The partial representation of the T-cell infiltration program in TAMs further implies that immune-associated transcriptional programs should not be interpreted as markers of a single immune lineage only. Rather, this program likely captures local immune activity, including T-cell presence and macrophage-associated signaling. Tumor cores generally displayed more spatially structured and locally enriched program activity, whereas invasive regions showed more diffuse and heterogeneous program organization. This suggests that the difference is not defined only by which cell types are present, but also by how cell populations engage stress, immune response, matrix-remodeling and metabolic programs within the local tissue niche. Thus, regional GBM biology emerges from the combined effects of cellular composition, spatial architecture and transcriptional reprogramming within shared lineages ^8, 9^. This has potential clinical implications because future diagnostic classification of GBM may benefit from incorporating functionally defined tumor regions based on spatial transcriptomic and metabolic states, rather than relying on histology alone. Such functional region mapping could help distinguish core-like, invasive-like, immune-rich or metabolically active tissue niches that may differ in treatment vulnerability.

The lipidomics data extends this by demonstrating that the core-edge transition is accompanied by broad, class-structured metabolic remodeling. Tumor cores were enriched in membrane and storage lipid classes, including PCs, PEs, ether-linked phospholipids, SMs, CEs and TGs, whereas invasive regions showed relative enrichment of Cer, GalCer and GlcCer. This is consistent with known roles of lipid metabolism in GBM growth, membrane synthesis, stress adaptation and therapy resistance ^17–19^. In particular, the enrichment of PCs and PEs in tumor regions supports increased membrane phospholipid demand in densely cellular and proliferative tumor areas, whereas CE and TG enrichment points toward lipid storage and buffering mechanisms that may protect tumor and immune cells from metabolic stress ^20, 21^. The invasive enrichment of ceramides could suggest a distinct lipid remodeling state at the margin, potentially reflecting membrane turnover, tissue remodeling, or altered lipid signaling in regions where tumor cells interact with the neural parenchyma. Importantly, lipid remodeling was not uniform across cell types. PCs were broadly tumor-enriched across malignant, stromal, and immune compartments, suggesting a core-associated membrane lipid state. In contrast, PEs, sphingolipids, and glycosphingolipids showed more species-and cell-type-dependent behavior, with both tumor-enriched and invasive-enriched patterns observed across different cellular types. This distinction is important because it argues against interpreting the differences in lipids as purely bulk. Instead, the same tissue region contains multiple lipid programs that are locally differentially associated with malignant, immune, and neural compartments. These findings are based upon previous MALDI-MSI studies of glioma lipid heterogeneity by assigning lipid changes to spatially defined cell identities and transcriptional states ^4, 5, 22^. In a translational context, such lipid-class and lipid-species patterns could eventually contribute to metabolic tissue signatures that distinguish core-like from invasive-like regions. If validated in larger cohorts, spatial lipid panels may support surgical margin assessment, imaging-based metabolic mapping, or biomarker development for infiltration state and regional aggressiveness.

One important conceptual advance of this work is the identification of spatial lipid-transcriptional coupling. Rather than treating transcriptomics and lipidomics as parallel layers, it was discovered that specific lipid classes and lipid species covariate with gene programs in the same spatial context. Tumor cores showed stronger and clearer lipid-gene coupling than invasive regions, indicating that transcriptional states in the core are more tightly linked to local lipid composition. This was particularly evident for lipid classes associated with membrane composition (PC, PE, PI) lipid storage (DG, TG), and lipid signaling (Cer). In contrast, the invasive region showed more heterogeneous and less strong coupling, suggesting that transcriptional and lipid states become partially stronger in a tumor rich tissue. The TAMs phagocytic program and CE relationship was especially showing. TAMs phagocytic activity spatially co-localized with CE abundance in discrete tissue regions containing TAMs. TAMs may phagocytose membrane debris, apoptotic material, myelin-derived lipids, and cholesterol-rich substrates in tumor regions, after which excess cholesterol can be esterified and stored as cholesteryl esters. This points to TAM-rich niches as local lipid-processing compartments. This adds a spatially resolved dimension to prior work linking cholesterol esterification and lipid storage to GBM growth and metabolic adaptation ^17, 49, 50^. These findings position TAMs not only as immune-regulatory cells, but also as metabolically active components of the tumor ecosystem. In this view, TAM lipid uptake, cholesterol processing and lipid storage may represent therapeutic vulnerabilities, either by directly disrupting macrophage lipid handling or by combining metabolic interventions with immune-modulating therapies. Importantly, these immune-associated modules should not be interpreted as complete functional immune-state annotations. The Xenium panel used here does not resolve CD8 T-cell identity or exhaustion states, and TAMs in GBM are known to span immunosuppressive, phagocytic, lipid-handling, and antigen-presentation-associated phenotypes. Therefore, the TAM phagocytic and T-cell infiltration programs are best interpreted as spatial immune-niche readouts rather than definitive classifications of macrophage activation or T-cell functional state. The lipid synthesis and PC relationship links transcriptional lipid activity to membrane phospholipid abundance, but this coupling was not uniform across the tissue. Regions with high lipid synthesis scores often corresponded with increased PC abundance, particularly in astrocyte-enriched tumor areas, supporting a local membrane remodeling state. However, regions with high PC signal but low lipid synthesis activity were also observed, indicating that PC abundance is not solely driven by local synthesis. Instead, PC levels could likely reflect multiple processes, including lipid uptake, membrane turnover and tissue composition. Thus, the spatially variable lipid synthesis and PC relationship implies that lipid synthesis explains part, but not all, of the membrane lipid landscape across the GBM tumor-edge axis ^21^. The T-cell infiltration and SM coupling was more heterogeneous, but it suggests that immune-associated niches may also carry distinct lipid environments. Sphingolipids are involved in membrane organization, signaling, and immune cell function, and their variable association with T-cell rich regions may reflect immune lipid microenvironments. This interpretation should remain cautious, but it highlights a potentially important area for future work: whether local lipid composition modulates immune cell localization, activation, or suppression in GBM ^14, 49^. Importantly, these spatial co-localization and correlation analyses identify coordinated molecular states, but they do not establish causality between transcriptional program activity and lipid abundance.

Finally, our exploratory MGMT analysis hints that the clinically relevant MGMT promoter methylation may be associated with differences in spatial lipid-transcriptional organization. MGMT methylation is most often interpreted as a predictive biomarker of TMZ benefit, based on the landmark finding that patients with MGMT^+^ glioblastoma derive greater benefit from TMZ treatment ^51, 52^. In our data, the strongest differences emerged at the level of gene-lipid coupling, particularly in the TAMs phagocytic program. This is important because it suggests that MGMT status may not primarily define whether a lipid class or gene program is globally increased or decreased, but rather how transcriptional states are metabolically implemented within specific cellular compartments. This raises the possibility that MGMT status may stratify patients not only by probability of response to chemotherapy, but also by immune-metabolic architecture and macrophage functional state within the tumor microenvironment.

MGMT^+^ tumors showed increased CE abundance and stronger tumor-associated TAM phagocytic program differences, accompanied by a distinct TAM lipid-gene coupling architecture. This pattern shows that MGMT^+^ tumors could contain macrophage lineages in which phagocytic, immune-regulatory and lipid handling programs are more tightly coordinated. In this context, CE enrichment may reflect increased uptake and processing of cholesterol-rich material by TAMs, potentially arising from tumor-cell membrane turnover or apoptotic debris. This interpretation is in line with the emerging view that MGMT promoter methylation captures more than DNA repair capacity alone ^53, 54^. Clinically, this suggests that treatment response may depend not only on tumor-cell DNA repair capacity, but also on the metabolic state of tumor-immune niches. In the future, MGMT-defined immune-metabolic differences could motivate combination strategies in which temozolomide-based therapy is paired with agents targeting macrophage activation, cholesterol handling or lipid storage pathways.

The TAMs-focused coupling analysis further indicates that TAMs may be important at which MGMT associated metabolic differences become visible. MGMT^+^ and MGMT^-^ tumors showed distinct patterns of coupling. With MGMT^+^ tumors showing stronger links between immune and phagocytic programs and lipid storage or membrane remodeling. These findings should be interpreted cautiously because the MGMT analysis is exploratory and limited by cohort size and therefore requires validation in larger patient cohorts. Nevertheless, it raises an important hypothesis: clinically defined GBM subgroups may differ not only in treatment sensitivity, but also in how tumor and immune cells coordinate transcriptional programs with lipid metabolism in space. If validated in larger cohorts, MGMT associated lipid and gene coupling could provide a framework for understanding why tumors with similar histological diagnosis, but different clinical biomarkers, may exhibit distinct metabolic microenvironments. More broadly, these results support the idea that spatial multimodal profiling can reveal clinically relevant tumor states that are not apparent from transcriptomic or lipidomic abundance measurements alone. This may ultimately support a more refined form of GBM subtyping in which molecular classification is complemented by spatial metabolic immune phenotypes, capturing how tumor cells, immune cells, and lipid states are organized within functional tissue niches.

Because the single-section MALDI-MSI-Xenium dataset is highly multidimensional, the present study follows a focused analytical trajectory: core-edge cellular composition, transcriptional programs, lipid remodeling, lipid-program coupling and exploratory MGMT-associated analysis. Other clinically or biologically relevant axes, including vascular niches, neuronal-tumor interactions, therapy-associated states, patient-specific metabolic programs, and additional molecular subgroups remain to be investigated. Our findings support a model in which the GBM core-edge axis is organized by coordinated changes in cellular composition, transcriptional state and lipid metabolism. By resolving these relationships within the same tissue section, this study provides a spatially grounded framework for understanding GBM metabolic heterogeneity and identifies lipid-gene program coupling as a potential organizing principle of tumor microenvironment biology. Combined, this data demonstrates that spatial lipid-transcriptomic profiling could help move GBM interpretation from static histological regions toward functional metabolic niches with potential diagnostic, prognostic and therapeutic relevance.

## Acknowledgments

The authors would like to thank Ruben Jacobs (Maastricht University) for performing the Xenium Spatial Transcriptomics experiment and Benjamin Balluff (Maastricht University) for helping with the python code. The authors would also like to thank the Department of Neurosurgery at KU Leuven for providing the GBM samples.

## Author contributions

T.F.E.H. contributed to the investigation, formal analysis, conceptualization, writing - review, and editing; G.B.E. contributed to software design and development, conceptualization and review; M.P.G.B. contributed to conceptualization and review; A.H. contributed to conceptualization and review; S.V. contributed to conceptualization and review; R.M.A.H. contributed to supervision, review and editing, funding acquisition; E.C. contributed to conceptualization, supervision, writing - review and editing, funding acquisition.

## Author information

### Data availability

Glioblastoma data cannot be shared directly, as this falls outside the scope of the approved ethical consents. Access to this data requires additional institutional review and adherence to the original patient consent.

### Code availability

We used ESCDAT Version P1 04-06-2025 for the overlay in this manuscript. The software packages such as R, Seurat and Python are publicly available and well documented. The Xenium to GeoJSON package and MALDI-MSI – Xenium SPT software ‘ESCDAT’ underlying these analysis are deposited at GitHub (https://github.com/M4i-Imaging-Mass-Spectrometry/MALDI-MSI---Spatial-Transcriptomics-Overlay).

### Additional Information

The authors declare no competing financial interest.

## Supporting information

SF1 – Patient level spatial maps; SF2 – Marker gene validation; SF3 – RNA UMAPs anatomical regions; SF4 – Cell type gene enrichment; SF5 – Uncorrected MALDI-MSI embedding; SF6 – Tumor core lipid-gene correlation; SF7 – Invasive zone lipid-gene correlation; SF8 – Spatial program proliferation; SF9 – Spatial program hypoxia; SF10 – Spatial program ECM; SF11 – Spatial program TAM phagocytic; SF12 – Spatial program T-cell infiltration; SF13 – Spatial program lipid synthesis; ST1 – Marker genes cell typing; ST2 – Patient metadata; ST3 – Patient cell information; ST4 – Marker genes gene programs; ST5 – Annotated lipid features.

## References

(1) Louis, D. N.; Perry, A.; Wesseling, P.; Brat, D. J.; Cree, I. A.; Figarella-Branger, D.; Hawkins, C.; Ng, H. K.; Pfister, S. M.; Reifenberger, G.;, et al. The 2021 WHO Classification of Tumors of the Central Nervous System: a summary. Neuro Oncol 2021, 23 (8), 1231–1251. DOI: 10.1093/neuonc/noab106 From NLM Medline.

(2) Sottoriva, A.; Spiteri, I.; Piccirillo, S. G.; Touloumis, A.; Collins, V. P.; Marioni, J. C.; Curtis, C.; Watts, C.; Tavare, S. Intratumor heterogeneity in human glioblastoma reflects cancer evolutionary dynamics. Proc Natl Acad Sci U S A 2013, 110 (10), 4009–4014. DOI: 10.1073/pnas.1219747110 From NLM Medline.

(3) Nomura, M.; Spitzer, A.; Johnson, K. C.; Garofano, L.; Nehar-Belaid, D.; Galili Darnell, N.; Greenwald, A. C.; Bussema, L.; Oh, Y. T.; Varn, F. S.;, et al. The multilayered transcriptional architecture of glioblastoma ecosystems. Nat Genet 2025, 57 (5), 1155–1167. DOI: 10.1038/s41588-025-02167-5 From NLM Medline.

(4) Roncevic, A.; Koruga, N.; Soldo Koruga, A.; Debeljak, Z.; Roncevic, R.; Turk, T.; Kretic, D.; Rotim, T.; Krivdic Dupan, Z.; Troha, D.;, et al. MALDI Imaging Mass Spectrometry of High-Grade Gliomas: A Review of Recent Progress and Future Perspective. Curr Issues Mol Biol 2023, 45 (2), 838–851. DOI: 10.3390/cimb45020055 From NLM PubMed-not-MEDLINE.

(5) Jha, D.; Blennow, K.; Zetterberg, H.; Savas, J. N.; Hanrieder, J. Spatial neurolipidomics-MALDI mass spectrometry imaging of lipids in brain pathologies. J Mass Spectrom 2024, 59 (3), e5008. DOI: 10.1002/jms.5008 From NLM Medline.

(6) Patel, A. P.; Tirosh, I.; Trombetta, J. J.; Shalek, A. K.; Gillespie, S. M.; Wakimoto, H.; Cahill, D. P.; Nahed, B. V.; Curry, W. T.; Martuza, R. L.;, et al. Single-cell RNA-seq highlights intratumoral heterogeneity in primary glioblastoma. Science 2014, 344 (6190), 1396–1401. DOI: 10.1126/science.1254257 From NLM Medline.

(7) Venkataramani, V.; Yang, Y.; Schubert, M. C.; Reyhan, E.; Tetzlaff, S. K.; Wissmann, N.; Botz, M.; Soyka, S. J.; Beretta, C. A.; Pramatarov, R. L.;, et al. Glioblastoma hijacks neuronal mechanisms for brain invasion. Cell 2022, 185 (16), 2899–2917 e2831. DOI: 10.1016/j.cell.2022.06.054 From NLM Medline.

(8) Hara, T.; Chanoch-Myers, R.; Mathewson, N. D.; Myskiw, C.; Atta, L.; Bussema, L.; Eichhorn, S. W.; Greenwald, A. C.; Kinker, G. S.; Rodman, C.;, et al. Interactions between cancer cells and immune cells drive transitions to mesenchymal-like states in glioblastoma. Cancer Cell 2021, 39 (6), 779–792 e711. DOI: 10.1016/j.ccell.2021.05.002 From NLM Medline.

(9) Wang, Q.; Hu, B.; Hu, X.; Kim, H.; Squatrito, M.; Scarpace, L.; deCarvalho, A. C.; Lyu, S.; Li, P.; Li, Y.;, et al. Tumor Evolution of Glioma-Intrinsic Gene Expression Subtypes Associates with Immunological Changes in the Microenvironment. Cancer Cell 2017, 32 (1), 42–56 e46. DOI: 10.1016/j.ccell.2017.06.003 From NLM Medline.

(10) Puchalski, R. B.; Shah, N.; Miller, J.; Dalley, R.; Nomura, S. R.; Yoon, J. G.; Smith, K. A.; Lankerovich, M.; Bertagnolli, D.; Bickley, K.;, et al. An anatomic transcriptional atlas of human glioblastoma. Science 2018, 360 (6389), 660–663. DOI: 10.1126/science.aaf2666 From NLM Medline.

(11) Wang, L.; Babikir, H.; Muller, S.; Yagnik, G.; Shamardani, K.; Catalan, F.; Kohanbash, G.; Alvarado, B.; Di Lullo, E.; Kriegstein, A.;, et al. The Phenotypes of Proliferating Glioblastoma Cells Reside on a Single Axis of Variation. Cancer Discov 2019, 9 (12), 1708–1719. DOI: 10.1158/2159-8290.CD-19-0329 From NLM Medline.

(12) Ruiz-Moreno, C.; Salas, S. M.; Samuelsson, E.; Minaeva, M.; Ibarra, I.; Grillo, M.; Brandner, S.; Roy, A.; Forsberg-Nilsson, K.; Kranendonk, M. E. G.;, et al. Charting the single-cell and spatial landscape of IDH-wild-type glioblastoma with GBmap. Neuro Oncol 2025, 27 (9), 2281–2295. DOI: 10.1093/neuonc/noaf113 From NLM Medline.

(13) Khan, S. M.; Wang, A. Z.; Desai, R. R.; McCornack, C. R.; Sun, R.; Dahiya, S. M.; Foltz, J. A.; Sherpa, N. D.; Leavitt, L.; West, T.;, et al. Mapping the spatial architecture of glioblastoma from core to edge delineates niche-specific tumor cell states and intercellular interactions. bioRxiv 2025. DOI: 10.1101/2025.04.04.647096 From NLM PubMed-not-MEDLINE.

(14) Karimi, E.; Yu, M. W.; Maritan, S. M.; Perus, L. J. M.; Rezanejad, M.; Sorin, M.; Dankner, M.; Fallah, P.; Dore, S.; Zuo, D.;, et al. Single-cell spatial immune landscapes of primary and metastatic brain tumours. Nature 2023, 614 (7948), 555–563. DOI: 10.1038/s41586-022-05680-3 From NLM Medline.

(15) Ren, Y.; Huang, Z.; Zhou, L.; Xiao, P.; Song, J.; He, P.; Xie, C.; Zhou, R.; Li, M.; Dong, X.;, et al. Spatial transcriptomics reveals niche-specific enrichment and vulnerabilities of radial glial stem-like cells in malignant gliomas. Nat Commun 2023, 14 (1), 1028. DOI: 10.1038/s41467-023-36707-6 From NLM Medline.

(16) Pai, B.; Ramos, S. I.; Cheng, W. S.; Joshi, T.; Ozen, E.; Kulumani Mahadevan, L. S.; Silva-Hurtado, T. J.; Price, G. A.; Tome-Garcia, J.; Nudelman, G.;, et al. Spatial Multiomics Defines a Shared Tumor Infiltrative Signature at the Resection Margin in High-Grade Gliomas. Cancer Res 2025, 85 (21), 4233–4250. DOI: 10.1158/0008-5472.CAN-24-4708 From NLM Medline.

(17) Geng, F.; Cheng, X.; Wu, X.; Yoo, J. Y.; Cheng, C.; Guo, J. Y.; Mo, X.; Ru, P.; Hurwitz, B.; Kim, S. H.;, et al. Inhibition of SOAT1 Suppresses Glioblastoma Growth via Blocking SREBP-1-Mediated Lipogenesis. Clin Cancer Res 2016, 22 (21), 5337–5348. DOI: 10.1158/1078-0432.CCR-15-2973 From NLM Medline.

(18) Miska, J.; Chandel, N. S. Targeting fatty acid metabolism in glioblastoma. J Clin Invest 2023, 133 (1). DOI: 10.1172/JCI163448 From NLM Medline.

(19) Xiao, M.; Xu, J.; Wang, W.; Zhang, B.; Liu, J.; Li, J.; Xu, H.; Zhao, Y.; Yu, X.; Shi, S. Functional significance of cholesterol metabolism in cancer: from threat to treatment. Exp Mol Med 2023, 55 (9), 1982–1995. DOI: 10.1038/s12276-023-01079-w From NLM Medline.

(20) Lu, L.; Zhang, Y.; Yang, Y.; Jin, M.; Ma, A.; Wang, X.; Zhao, Q.; Zhang, X.; Zheng, J.; Zheng, X. Lipid metabolism: the potential therapeutic targets in glioblastoma. Cell Death Discov 2025, 11 (1), 107. DOI: 10.1038/s41420-025-02390-3 From NLM PubMed-not-MEDLINE.

(21) Glowacka, K.; Ibanez, S.; Renoult, O.; Vermonden, P.; Giolito, M. V.; Ozkan, K.; Degavre, C.; Aubert, L.; Guilbaud, C.; Laloux-Morris, F.;, et al. Acid-exposed and hypoxic cancer cells do not overlap but are interdependent for unsaturated fatty acid resources. Nat Commun 2024, 15 (1), 10107. DOI: 10.1038/s41467-024-54435-3 From NLM Medline.

(22) Wood, J.; Smith, S. J.; Castellanos-Uribe, M.; Lourdusamy, A.; May, S. T.; Barrett, D. A.; Grundy, R. G.; Kim, D. H.; Rahman, R. Metabolomic characterisation of the glioblastoma invasive margin reveals a region-specific signature. Heliyon 2025, 11 (1), e41309. DOI: 10.1016/j.heliyon.2024.e41309 From NLM PubMed-not-MEDLINE.

(23) Sekera, E. R.; Rosas, L.; Holbrook, J. H.; Angeles-Lopez, Q. D.; Khaliullin, T.; Rojas, M.; Mora, A. L.; Hummon, A. B. Single Cell MALDI-MSI Analysis of Lipids and Proteins within a Replicative Senescence Fibroblast Model. J Am Soc Mass Spectrom 2024, 35 (12), 2815–2823. DOI: 10.1021/jasms.4c00095 From NLM Medline.

(24) Krestensen, K. K.; Heeren, R. M. A.; Balluff, B. State-of-the-art mass spectrometry imaging applications in biomedical research. Analyst 2023, 148 (24), 6161–6187. DOI: 10.1039/d3an01495a From NLM Medline.

(25) Alexandrov, T.; Saez-Rodriguez, J.; Saka, S. K. Enablers and challenges of spatial omics, a melting pot of technologies. Mol Syst Biol 2023, 19 (11), e10571. DOI: 10.15252/msb.202110571 From NLM Medline.

(26) Hendriks, T. F. E.; Eijkel, G. B.; Visvikis, T.; Balluff, B.; Heeren, R. M. A.; Cuypers, E. One section, two worlds: single-cell integration of MALDI-MSI and spatial transcriptomics on the same single tissue section. Sci Rep 2025, 15 (1), 42660. DOI: 10.1038/s41598-025-26735-1 From NLM Medline.

(27) Breitkopf, S. B.; Ricoult, S. J. H.; Yuan, M.; Xu, Y.; Peake, D. A.; Manning, B. D.; Asara, J. M. A relative quantitative positive/negative ion switching method for untargeted lipidomics via high resolution LC-MS/MS from any biological source. Metabolomics 2017, 13 (3). DOI: 10.1007/s11306-016-1157-8 From NLM PubMed-not-MEDLINE.

(28) Scupakova, K.; Dewez, F.; Walch, A. K.; Heeren, R. M. A.; Balluff, B. Morphometric Cell Classification for Single-Cell MALDI-Mass Spectrometry Imaging. Angew Chem Int Ed Engl 2020, 59 (40), 17447–17450. DOI: 10.1002/anie.202007315 From NLM Medline.

(29) Butler, A.; Hoffman, P.; Smibert, P.; Papalexi, E.; Satija, R. Integrating single-cell transcriptomic data across different conditions, technologies, and species. Nat Biotechnol 2018, 36 (5), 411–420. DOI: 10.1038/nbt.4096 From NLM Medline.

(30) Hao, Y.; Stuart, T.; Kowalski, M. H.; Choudhary, S.; Hoffman, P.; Hartman, A.; Srivastava, A.; Molla, G.; Madad, S.; Fernandez-Granda, C.;, et al. Dictionary learning for integrative, multimodal and scalable single-cell analysis. Nat Biotechnol 2024, 42 (2), 293–304. DOI: 10.1038/s41587-023-01767-y From NLM Medline.

(31) Satija, R.; Farrell, J. A.; Gennert, D.; Schier, A. F.; Regev, A. Spatial reconstruction of single-cell gene expression data. Nat Biotechnol 2015, 33 (5), 495–502. DOI: 10.1038/nbt.3192 From NLM Medline.

(32) Li, J.; Wang, P.; Wang, L. Y.; Wu, Y.; Wang, J.; Yu, D.; Chen, Z.; Shi, H.; Yin, S. Redistribution of the astrocyte phenotypes in the medial vestibular nuclei after unilateral labyrinthectomy. Front Neurosci 2023, 17, 1146147. DOI: 10.3389/fnins.2023.1146147 From NLM PubMed-not-MEDLINE.

(33) Yang, Y.; Chu, L.; Zeng, Z.; Xu, S.; Yang, H.; Zhang, X.; Jia, J.; Long, N.; Hu, Y.; Liu, J. Four specific biomarkers associated with the progression of glioblastoma multiforme in older adults identified using weighted gene co-expression network analysis. Bioengineered 2021, 12 (1), 6643–6654. DOI: 10.1080/21655979.2021.1975980 From NLM Medline.

(34) Amin, M.; Mays, M.; Polston, D.; Flanagan, E. P.; Prayson, R.; Kunchok, A. Myelin oligodendrocyte glycoprotein (MOG) antibodies in a patient with glioblastoma: Red flags for false positivity. J Neuroimmunol 2021, 361, 577743. DOI: 10.1016/j.jneuroim.2021.577743 From NLM Medline.

(35) Ye, D.; Wang, Q.; Yang, Y.; Chen, B.; Zhang, F.; Wang, Z.; Luan, Z. Identifying Genes that Affect Differentiation of Human Neural Stem Cells and Myelination of Mature Oligodendrocytes. Cell Mol Neurobiol 2023, 43 (5), 2337–2358. DOI: 10.1007/s10571-022-01313-5 From NLM Medline.

(36) Hayashida, S.; Masaki, K.; Suzuki, S. O.; Yamasaki, R.; Watanabe, M.; Koyama, S.; Isobe, N.; Matsushita, T.; Takahashi, K.; Tabira, T.;, et al. Distinct microglial and macrophage distribution patterns in the concentric and lamellar lesions in Balo’s disease and neuromyelitis optica spectrum disorders. Brain Pathol 2020, 30 (6), 1144–1157. DOI: 10.1111/bpa.12898 From NLM Medline.

(37) Mecca, C.; Giambanco, I.; Donato, R.; Arcuri, C. Microglia and Aging: The Role of the TREM2-DAP12 and CX3CL1-CX3CR1 Axes. Int J Mol Sci 2018, 19 (1). DOI: 10.3390/ijms19010318 From NLM Medline.

(38) Ravi, V. M.; Neidert, N.; Will, P.; Joseph, K.; Maier, J. P.; Kuckelhaus, J.; Vollmer, L.; Goeldner, J. M.; Behringer, S. P.; Scherer, F.;, et al. T-cell dysfunction in the glioblastoma microenvironment is mediated by myeloid cells releasing interleukin-10. Nat Commun 2022, 13 (1), 925. DOI: 10.1038/s41467-022-28523-1 From NLM Medline.

(39) Lichtenberger, B. M.; Tan, P. K.; Niederleithner, H.; Ferrara, N.; Petzelbauer, P.; Sibilia, M. Autocrine VEGF signaling synergizes with EGFR in tumor cells to promote epithelial cancer development. Cell 2010, 140 (2), 268–279. DOI: 10.1016/j.cell.2009.12.046 From NLM Medline.

(40) Tasic, B.; Yao, Z.; Graybuck, L. T.; Smith, K. A.; Nguyen, T. N.; Bertagnolli, D.; Goldy, J.; Garren, E.; Economo, M. N.; Viswanathan, S.;, et al. Shared and distinct transcriptomic cell types across neocortical areas. Nature 2018, 563 (7729), 72–78. DOI: 10.1038/s41586-018-0654-5 From NLM Medline.

(41) Kim, M. H.; Radaelli, C.; Thomsen, E. R.; Monet, D.; Chartrand, T.; Jorstad, N. L.; Mahoney, J. T.; Taormina, M. J.; Long, B.; Baker, K.;, et al. Target cell-specific synaptic dynamics of excitatory to inhibitory neuron connections in supragranular layers of human neocortex. Elife 2023, 12. DOI: 10.7554/eLife.81863 From NLM Medline.

(42) Ooki, A.; Dinalankara, W.; Marchionni, L.; Tsay, J. J.; Goparaju, C.; Maleki, Z.; Rom, W. N.; Pass, H. I.; Hoque, M. O. Epigenetically regulated PAX6 drives cancer cells toward a stem-like state via GLI-SOX2 signaling axis in lung adenocarcinoma. Oncogene 2018, 37 (45), 5967–5981. DOI: 10.1038/s41388-018-0373-2 From NLM Medline.

(43) Krepela, E.; Vanickova, Z.; Hrabal, P.; Zubal, M.; Chmielova, B.; Balaziova, E.; Vymola, P.; Matrasova, I.; Busek, P.; Sedo, A. Regulation of Fibroblast Activation Protein by Transforming Growth Factor Beta-1 in Glioblastoma Microenvironment. Int J Mol Sci 2021, 22 (3). DOI: 10.3390/ijms22031046 From NLM Medline.

(44) Li, Y.; Xu, X.; Wang, X.; Zhang, C.; Hu, A.; Li, Y. MGST1 Expression Is Associated with Poor Prognosis, Enhancing the Wnt/beta-Catenin Pathway via Regulating AKT and Inhibiting Ferroptosis in Gastric Cancer. ACS Omega 2023, 8 (26), 23683–23694. DOI: 10.1021/acsomega.3c01782 From NLM PubMed-not-MEDLINE.

(45) Balluff, B.; Hopf, C.; Porta Siegel, T.; Grabsch, H. I.; Heeren, R. M. A. Batch Effects in MALDI Mass Spectrometry Imaging. J Am Soc Mass Spectrom 2021, 32 (3), 628–635. DOI: 10.1021/jasms.0c00393 From NLM PubMed-not-MEDLINE.

(46) Sparre, A. A.; Jensen, O. N. Batch correction for large-scale mass spectrometry imaging experiments. Bioinformatics 2026, 42 (6). DOI: 10.1093/bioinformatics/btag360 From NLM Medline.

(47) Neftel, C.; Laffy, J.; Filbin, M. G.; Hara, T.; Shore, M. E.; Rahme, G. J.; Richman, A. R.; Silverbush, D.; Shaw, M. L.; Hebert, C. M.;, et al. An Integrative Model of Cellular States, Plasticity, and Genetics for Glioblastoma. Cell 2019, 178 (4), 835–849 e821. DOI: 10.1016/j.cell.2019.06.024 From NLM Medline.

(48) Greenwald, A. C.; Darnell, N. G.; Hoefflin, R.; Simkin, D.; Mount, C. W.; Gonzalez Castro, L. N.; Harnik, Y.; Dumont, S.; Hirsch, D.; Nomura, M.;, et al. Integrative spatial analysis reveals a multi-layered organization of glioblastoma. Cell 2024, 187 (10), 2485–2501 e2426. DOI: 10.1016/j.cell.2024.03.029 From NLM Medline.

(49) Pombo Antunes, A. R.; Scheyltjens, I.; Lodi, F.; Messiaen, J.; Antoranz, A.; Duerinck, J.; Kancheva, D.; Martens, L.; De Vlaminck, K.; Van Hove, H.;, et al. Single-cell profiling of myeloid cells in glioblastoma across species and disease stage reveals macrophage competition and specialization. Nat Neurosci 2021, 24 (4), 595–610. DOI: 10.1038/s41593-020-00789-y From NLM Medline.

(50) Kloosterman, D. J.; Erbani, J.; Boon, M.; Farber, M.; Handgraaf, S. M.; Ando-Kuri, M.; Sanchez-Lopez, E.; Fontein, B.; Mertz, M.; Nieuwland, M.;, et al. Macrophage-mediated myelin recycling fuels brain cancer malignancy. Cell 2024, 187 (19), 5336–5356 e5330. DOI: 10.1016/j.cell.2024.07.030 From NLM Medline.

(51) Hegi, M. E.; Diserens, A. C.; Gorlia, T.; Hamou, M. F.; de Tribolet, N.; Weller, M.; Kros, J. M.; Hainfellner, J. A.; Mason, W.; Mariani, L.;, et al. MGMT gene silencing and benefit from temozolomide in glioblastoma. N Engl J Med 2005, 352 (10), 997–1003. DOI: 10.1056/NEJMoa043331 From NLM Medline.

(52) Brandner, S.; McAleenan, A.; Kelly, C.; Spiga, F.; Cheng, H. Y.; Dawson, S.; Schmidt, L.; Faulkner, C. L.; Wragg, C.; Jefferies, S.;, et al. MGMT promoter methylation testing to predict overall survival in people with glioblastoma treated with temozolomide: a comprehensive meta-analysis based on a Cochrane Systematic Review. Neuro Oncol 2021, 23 (9), 1457–1469. DOI: 10.1093/neuonc/noab105 From NLM Medline.

(53) Chen, X.; Sun, J.; Li, Y.; Jiang, W.; Li, Z.; Mao, J.; Zhou, L.; Chen, S.; Tan, G. Proteomic and metabolomic analyses illustrate the mechanisms of expression of the O(6)-methylguanine-DNA methyltransferase gene in glioblastoma. CNS Neurosci Ther 2024, 30 (2), e14415. DOI: 10.1111/cns.14415 From NLM Medline.

(54) Kushihara, Y.; Tanaka, S.; Kobayashi, Y.; Nagaoka, K.; Kikuchi, M.; Nejo, T.; Yamazawa, E.; Nambu, S.; Kugasawa, K.; Takami, H.;, et al. Glioblastoma with high O6-methyl-guanine DNA methyltransferase expression are more immunologically active than tumors with low MGMT expression. Front Immunol 2024, 15, 1328375. DOI: 10.3389/fimmu.2024.1328375 From NLM Medline.

