## Supplemental information for "Spatial Single Cell Lipid-Transcriptomic Coupling Reveals Metabolic Niches in Glioblastoma"

### Cell-Type Lipidomics and Spatial Transcriptomics Maps the Core and Invasive Edge Transition in Glioblastoma

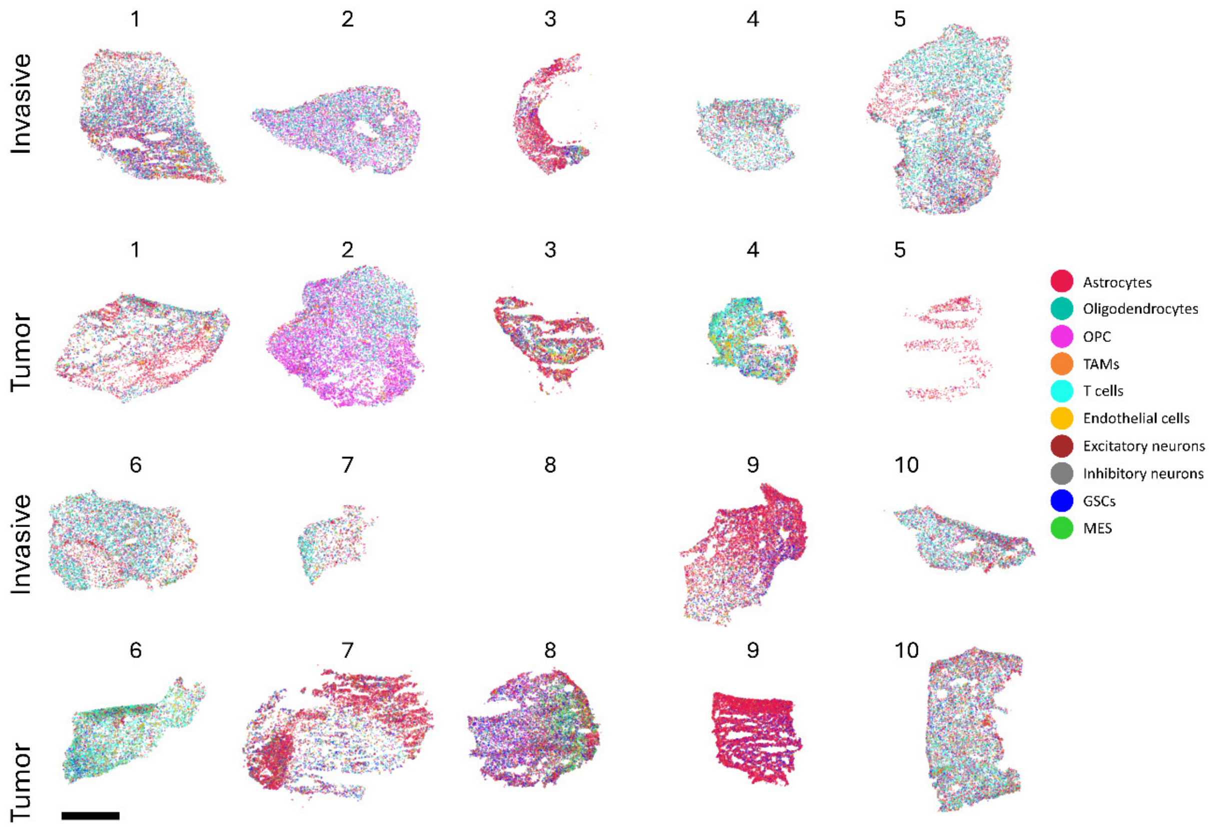


**Supplementary Figure S1. Patient-level spatial maps of annotated cell types across tumor-core and invasive-region sections**. Spatial Xenium cell-type maps are shown for all retained glioblastoma sections included in the study. Patient-matched invasive-region sections are displayed above their corresponding tumor-core sections where available. Each point represents a single segmented cell colored by annotation. One invasive-region section was excluded from downstream analysis, resulting in 19 retained sections. Scale bar indicates 1 mm.

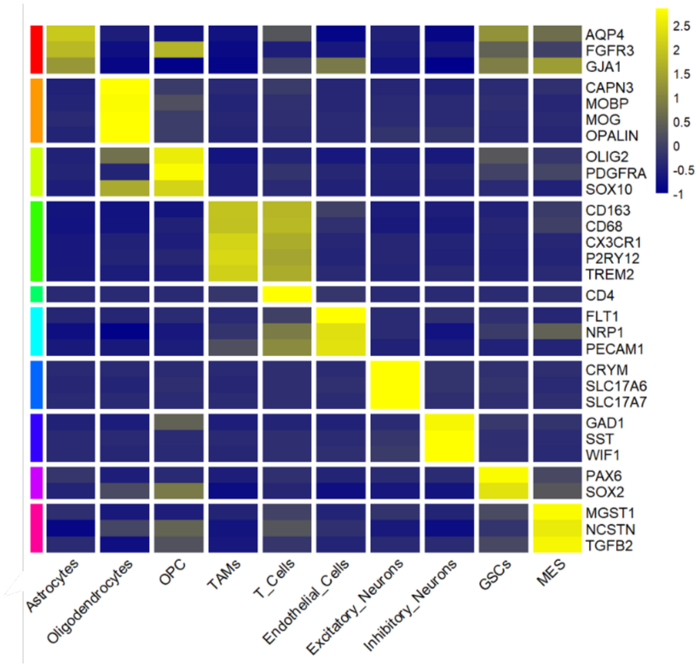


**Supplementary Figure S2. Marker-gene validation of Xenium-derived cell-type annotations.** Heatmap showing scaled expression of canonical marker genes across annotated cell types. Rows represent marker genes grouped by expected cellular identity, and columns represent annotated cell populations. Yellow indicates higher scaled expression, blue indicates lower expression.

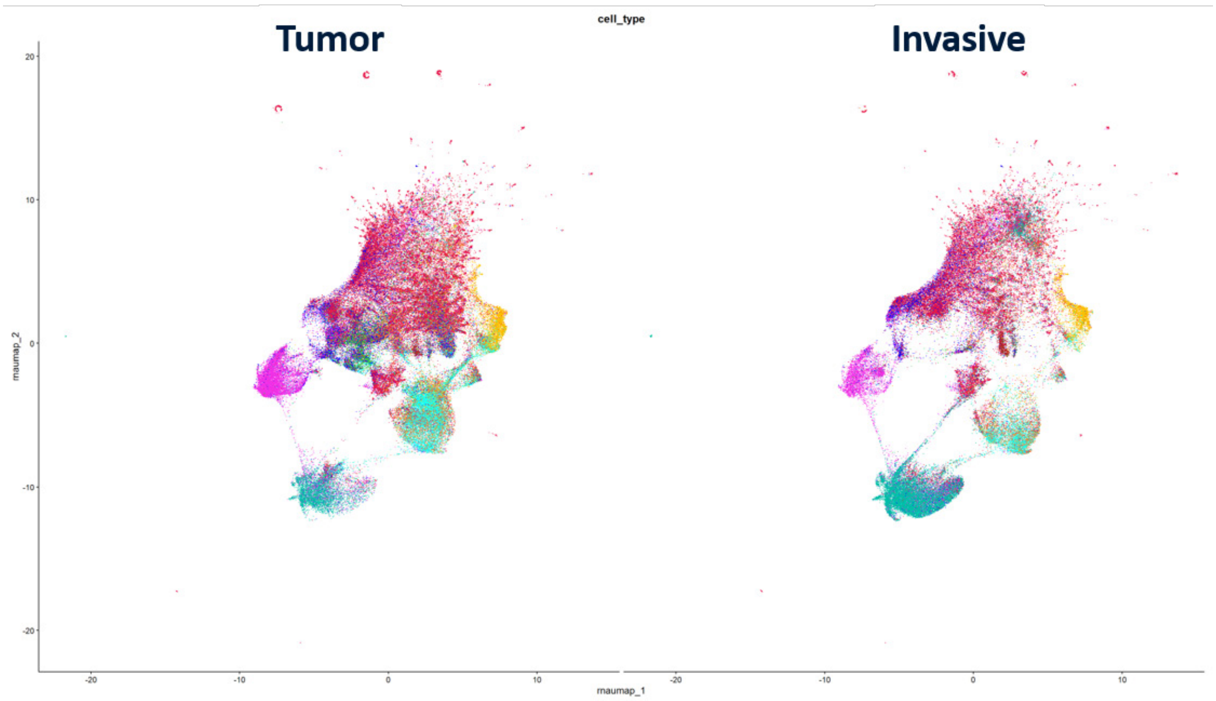


**Supplementary Figure S3. Joint RNA UMAP embedding of annotated cell types split by anatomical region.** UMAP embedding of Xenium RNA profiles from tumor-core and invasive-region sections, colored by cell-type annotation. The same UMAP coordinate space is shown for both regions, enabling direct comparison of cell-state occupancy across the tumor core - invasive edge axis. Major annotated cell populations are recovered in both tumor-core and invasive-region samples, indicating that the two compartments share a common transcriptional state space. Regional differences are reflected primarily by altered relative occupancy of shared cell-type neighborhoods.

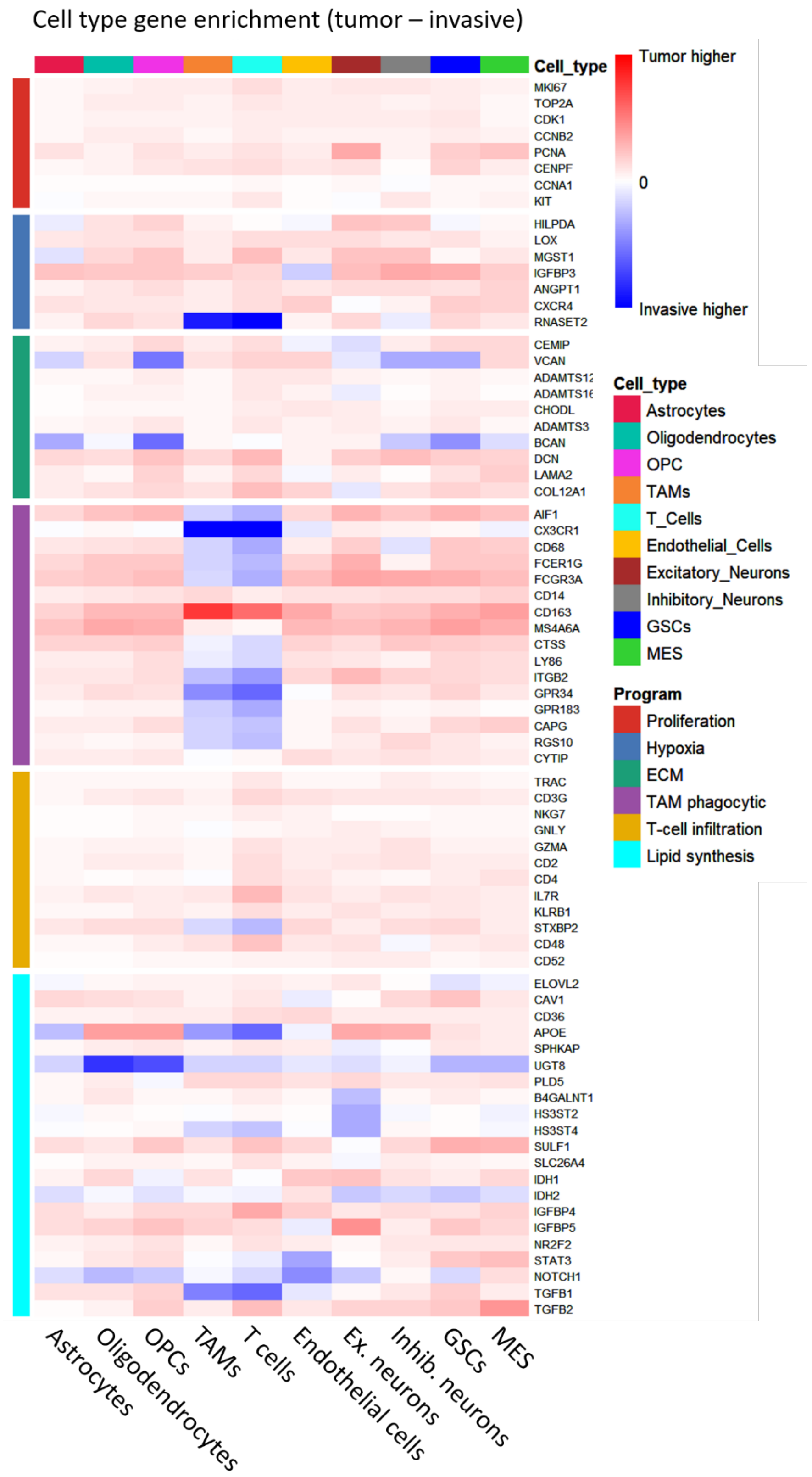


**Supplementary Figure S4. Heatmap showing tumor-core versus invasive-region differences in the expression of individual program-defining genes across annotated cell types.** Genes grouped by transcriptional program: proliferation, hypoxia, ECM remodeling, TAM phagocytic/lipid-handling activity, T-cell infiltration and lipid synthesis. Annotated cell types. (Red) higher expression in tumor-core samples. (Blue) indicates higher expression in invasive-region samples, and white indicates little regional differenceThis gene-level view complements the program-level summary in Figure 2C by showing that regional transcriptional differences are structured but not uniform across all genes within each program.

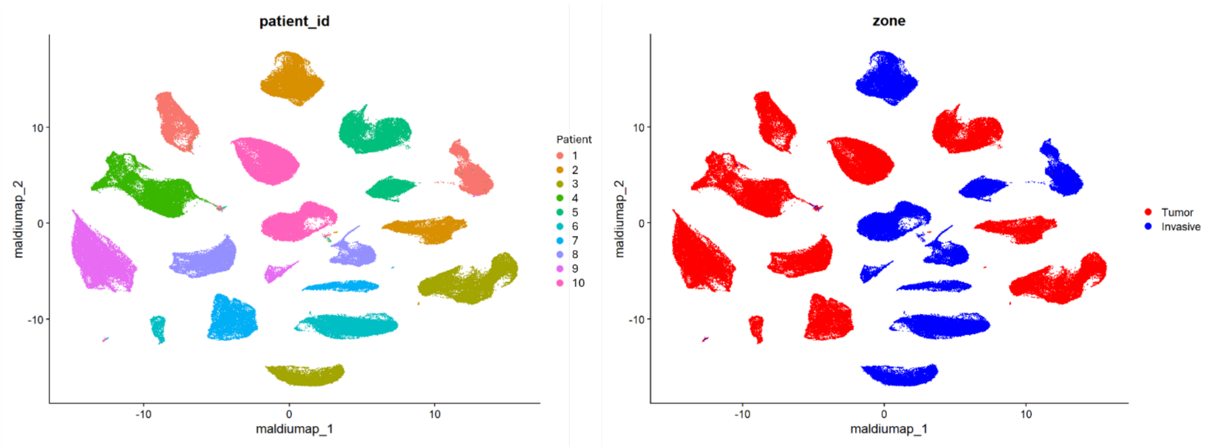


**Supplementary Figure S5. Uncorrected MALDI-MSI lipid embedding shows patient-associated batch structure.** UMAP embedding of MALDI-MSI lipid features prior to Harmony-based correction, colored by patient identity (left) and anatomical region (right). The uncorrected embedding clusters primarily by patient, indicating strong inter-patient variability in the raw lipid feature space.

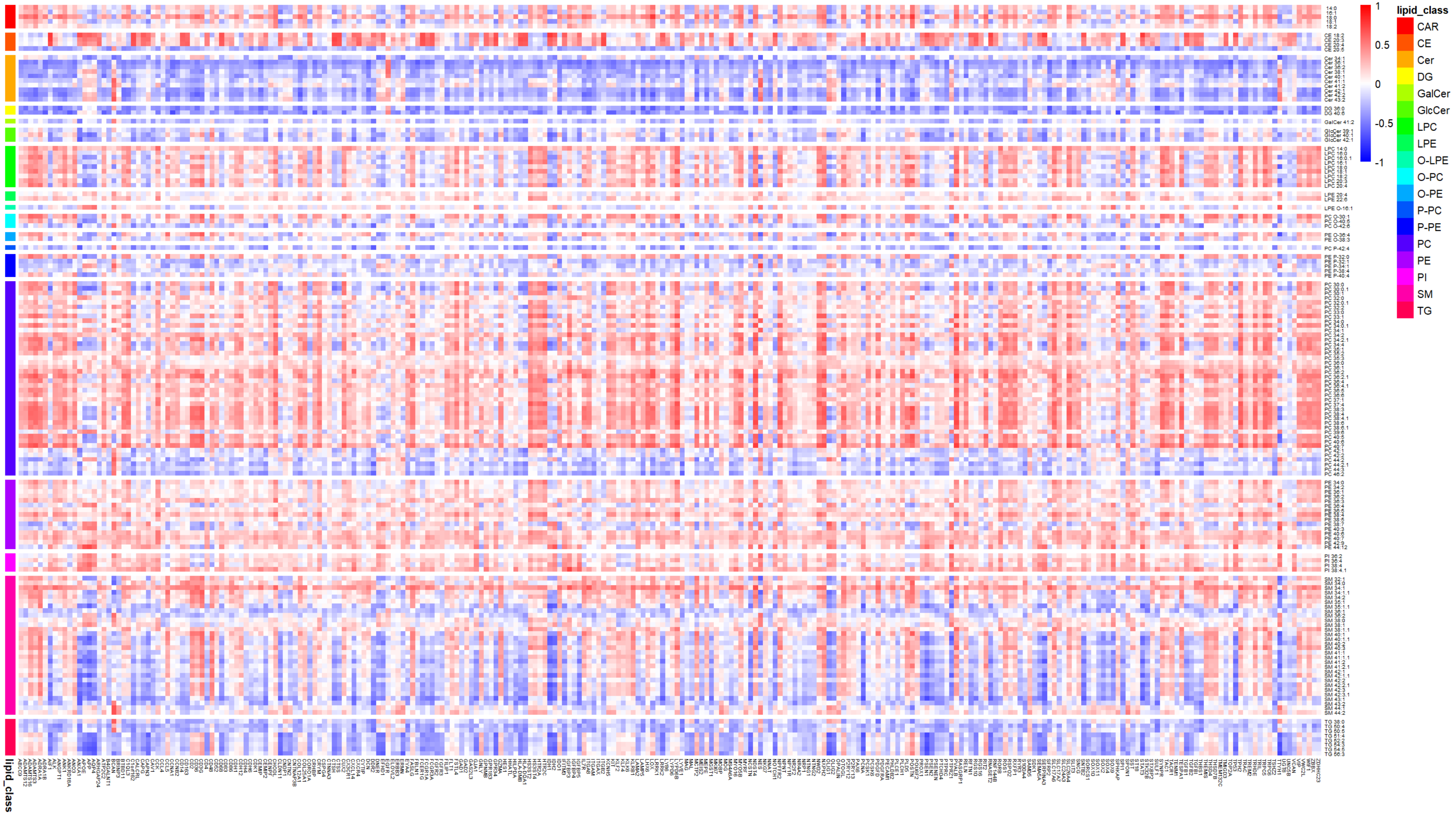

**Supplementary Figure S6. Tumor-core lipid-gene correlation matrix across annotated lipid species and Xenium genes.** Heatmap showing Spearman correlations between identified MALDI-MSI lipid species and all Xenium genes in tumor-core samples. Red indicates positive lipid-gene correlation, blue indicates negative correlation, and white indicates weak or no correlation. The tumor-core matrix shows structured lipid-class-dependent correlation patterns.

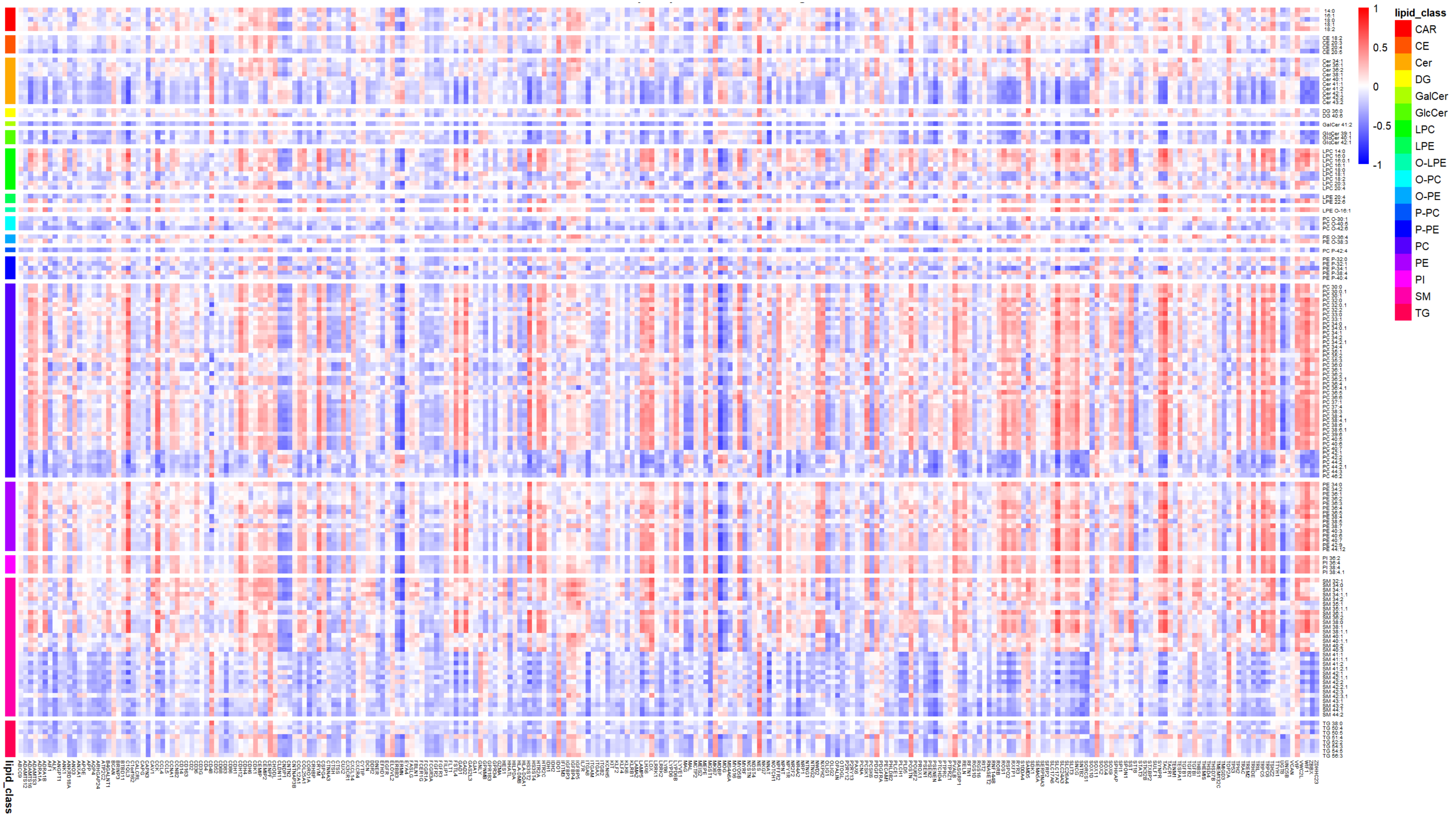


**Supplementary Figure S7.** Invasive-region lipid-gene correlation matrix across annotated lipid species and Xenium genes. Heatmap showing Spearman correlations between identified MALDI-MSI lipid species and all measured Xenium genes in invasive-region samples. Red indicates positive lipid-gene correlation, blue indicates negative correlation, and white indicates weak or no correlation. Compared with tumor-core regions, invasive samples show more heterogeneous and less block-like lipid-gene correlation structure.

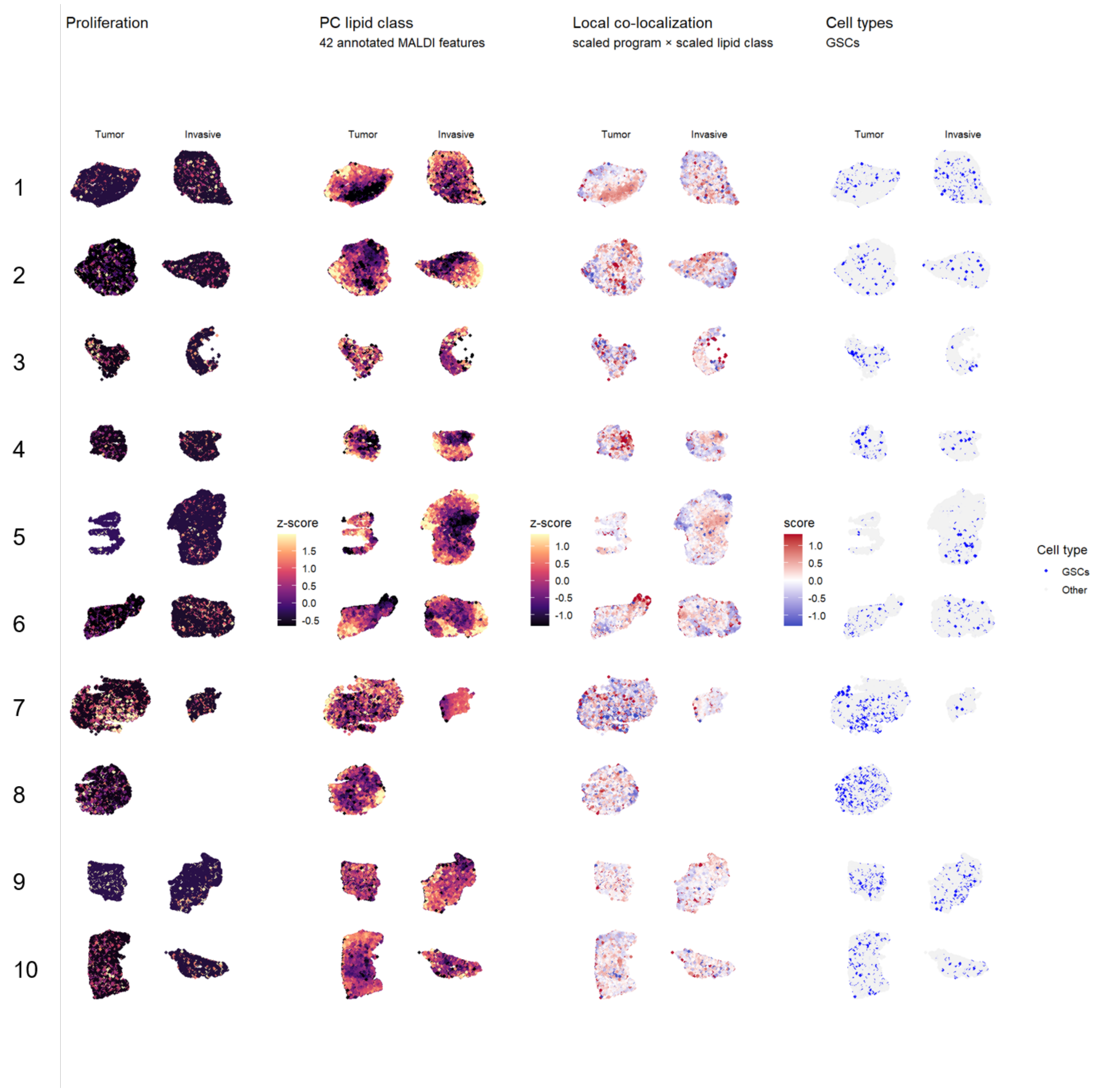


**Supplementary figure S8. Spatial program proliferation and PC coupling across matched tumor-core and invasive-region sections.** Spatial maps showing proliferation program scores, phosphatidylcholine (PC) lipid-class abundance, local co-localization, and GSC annotations. PC abundance was calculated from 42 annotated MALDI-MSI features. Program and lipid maps are shown as z-scores. Local co-localization was calculated as the product of scaled proliferation program score and scaled PC abundance; red indicates local co-enrichment, blue indicates local discordance, and white indicates weak or no co-localization.

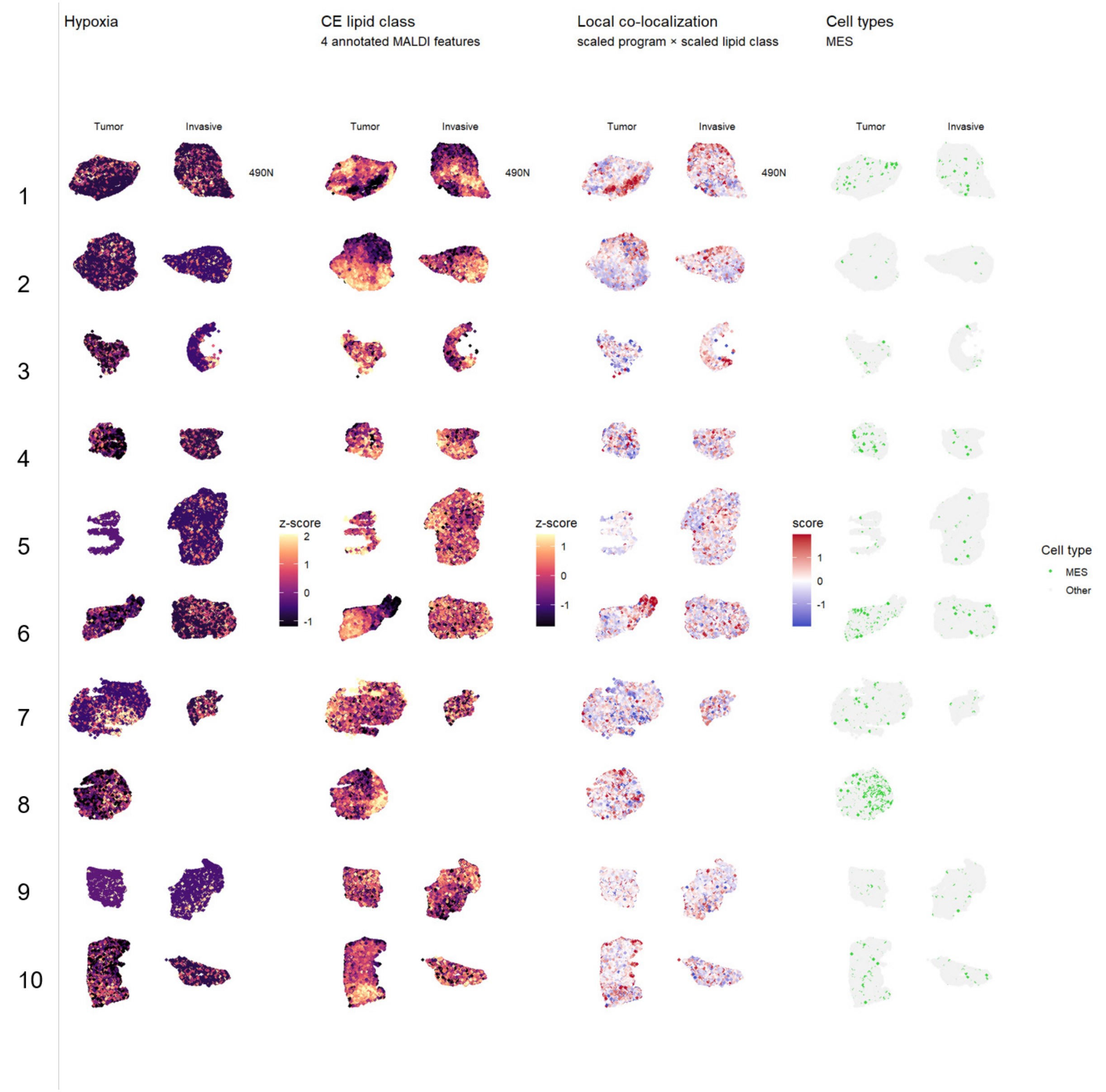


**Supplementary figure S9. Spatial hypoxia program and cholesteryl ester coupling across matched tumor-core and invasive-region sections.** Spatial maps showing hypoxia-associated program scores, cholesteryl ester (CE) lipid-class abundance, local co-localization, and MES-like cell annotations across matched tumor-core and invasive-region sections. CE abundance was calculated from 4 annotated MALDI-MSI features. Program and lipid maps are shown as z-scores. Local co-localization was calculated as the product of scaled hypoxia program score and scaled CE abundance; red indicates local co-enrichment, blue indicates local discordance, and white indicates weak or no co-localization.

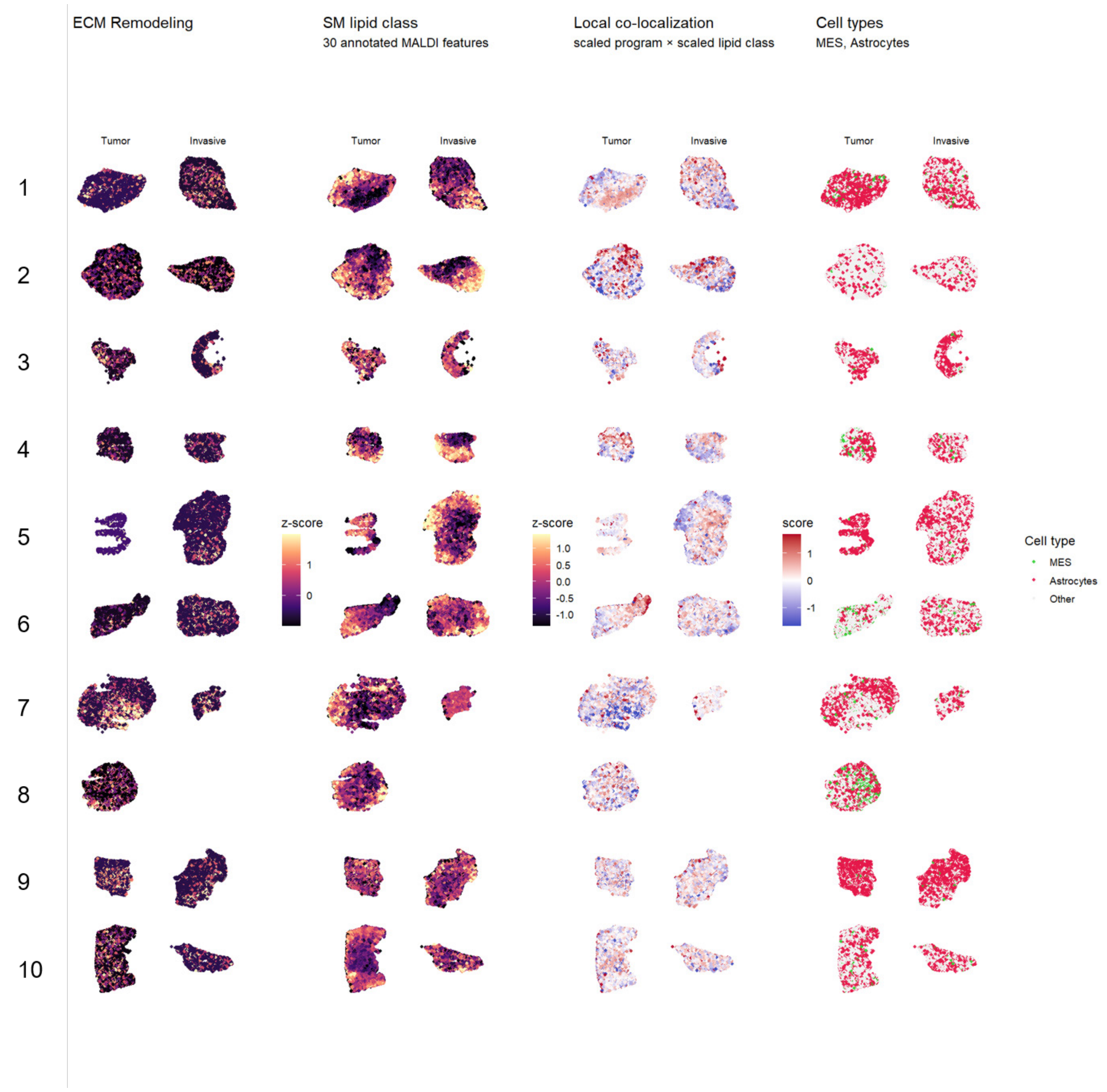


**Supplementary figure S10. Spatial ECM remodeling program and sphingomyelin coupling across matched tumor-core and invasive-region sections.** Spatial maps showing ECM remodeling program scores, sphingomyelin (SM) lipid-class abundance, local co-localization, and MES-like/astrocyte annotations across matched tumor-core and invasive-region sections. SM abundance was calculated from 30 annotated MALDI-MSI features. Program and lipid maps are shown as z-scores. Local co-localization was calculated as the product of scaled ECM remodeling program score and scaled SM abundance; red indicates local co-enrichment, blue indicates local discordance, and white indicates weak or no co-localization.

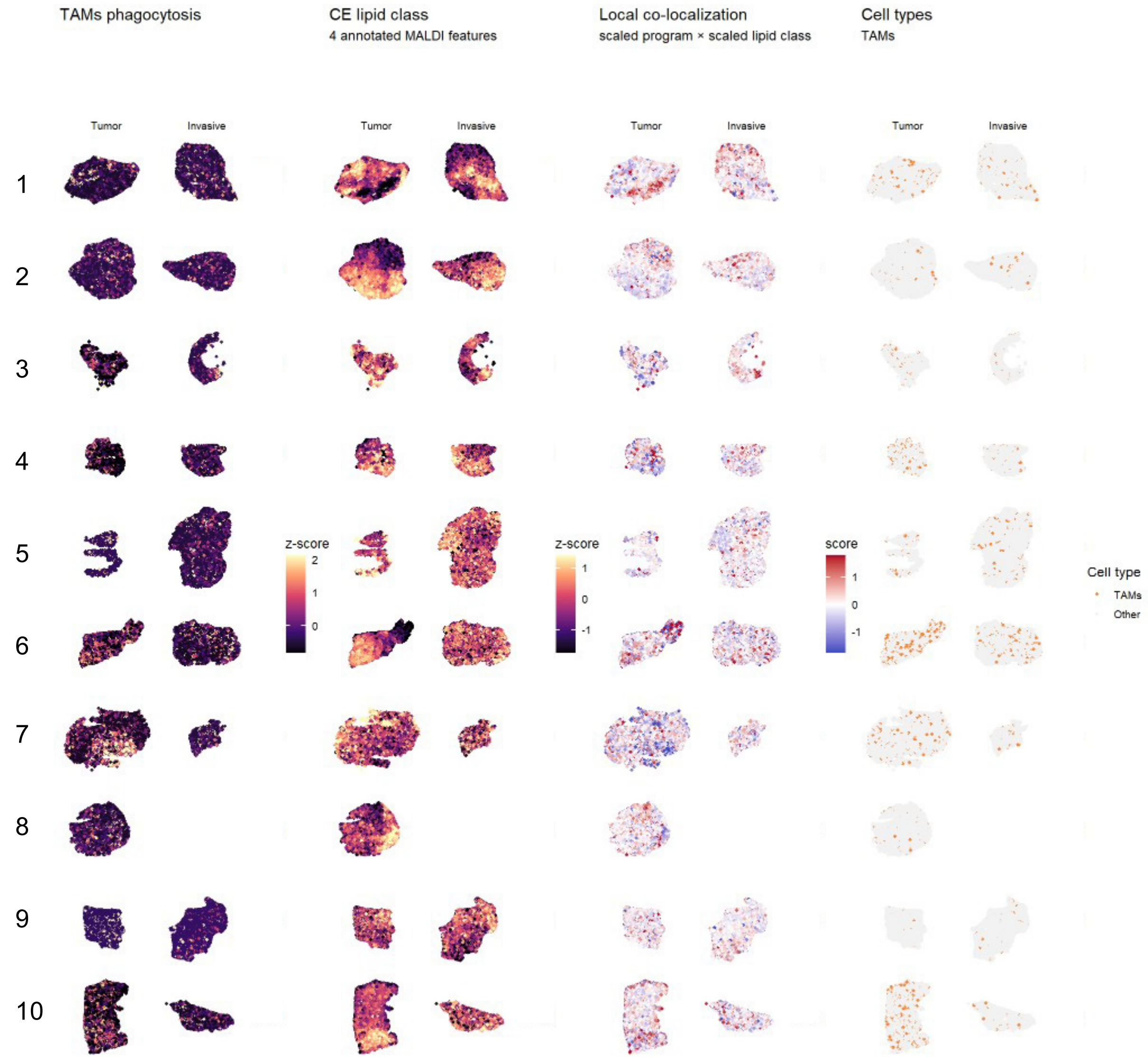


**Supplementary Figure S11. Spatial TAM phagocytic program and cholesteryl ester coupling across matched tumor-core and invasive-region sections.** Spatial maps showing TAM phagocytic program scores, cholesteryl ester (CE) lipid-class abundance, local co-localization, and TAM annotations across matched tumor-core and invasive-region sections. CE abundance was calculated from 4 annotated MALDI-MSI features. Program and lipid maps are shown as z-scores. Local co-localization was calculated as the product of scaled TAM phagocytic program score and scaled CE abundance; red indicates local co-enrichment, blue indicates local discordance, and white indicates weak or no co-localization.

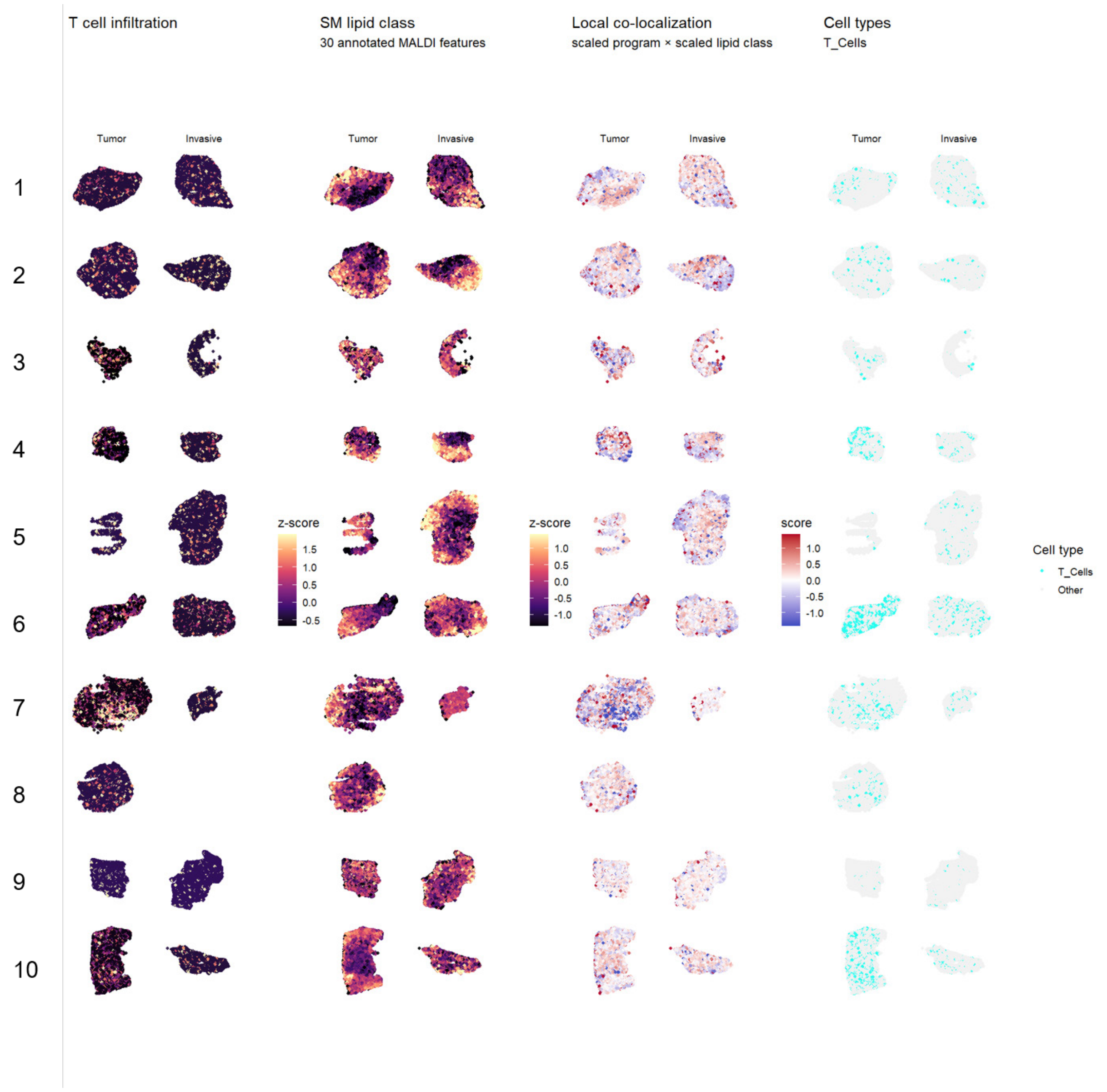


**Supplementary Figure S12. Spatial T-cell infiltration program and sphingomyelin coupling across matched tumor-core and invasive-region sections.** Spatial maps showing T-cell infiltration program scores, sphingomyelin (SM) lipid-class abundance, local co-localization, and T-cell annotations across matched tumor-core and invasive-region sections. SM abundance was calculated from 30 annotated MALDI-MSI features. Program and lipid maps are shown as z-scores. Local co-localization was calculated as the product of scaled T-cell infiltration program score and scaled SM abundance; red indicates local co-enrichment, blue indicates local discordance, and white indicates weak or no co-localization.

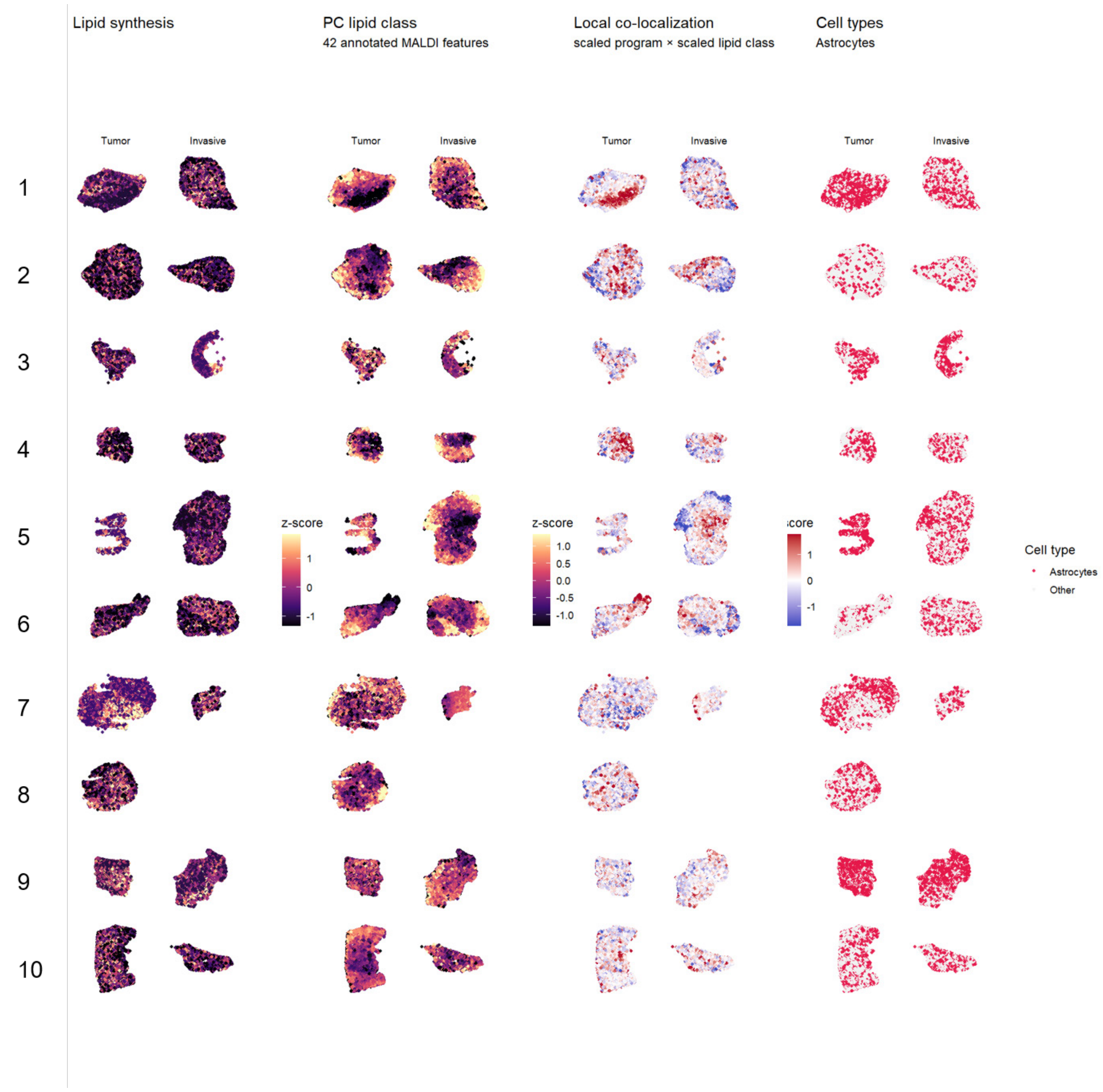


**Supplementary Figure S13. Spatial lipid synthesis program and phosphatidylcholine coupling across matched tumor-core and invasive-region sections.** Spatial maps showing lipid synthesis program scores, phosphatidylcholine (PC) lipid-class abundance, local co-localization, and astrocyte annotations across matched tumor-core and invasive-region sections. PC abundance was calculated from 42 annotated MALDI-MSI features. Program and lipid maps are shown as z-scores. Local co-localization was calculated as the product of scaled lipid synthesis program score and scaled PC abundance; red indicates local co-enrichment, blue indicates local discordance, and white indicates weak or no co-localization. Cell-type maps show astrocytes compared with all other annotated cells.

**Supplementary Table S1. Marker genes used for cell-type annotation.** The table lists the canonical marker genes used to support annotation of astrocytes, endothelial cells, excitatory neurons, GSCs, inhibitory neurons, MES-like cells, OPCs, oligodendrocytes, TAMs, and T cells.

| Cell type | Marker list |
|---|---|
| Astrocytes | AQP4, GJA1, FGFR3 |
| Endothelial cells | PECAM1, FLT1, NRP1 |
| Excitatory neurons | CRYM, SLC17A7, SLC17A6 |
| Glioblastoma stem cells (GSCs) | PAX6, SOX2 |
| Inhibitory neurons | GAD1, WIF1, SST |
| Mesenchymal cells (MES) | TGFB2, MGST1, NCSTN |
| Oligodendrocyte progenitor cells (OPCs) | SOX10, OLIG2, PDGFRA |
| Oligodendrocytes | CAPN3, MOBP, MOG, OPALIN |
| Tumor associated macrophages and microglia (TAMs) | CD163, CD68. CX3CR1. P2RY12, TREM2 |
| T cells | CD4 |

**Supplementary Table S2. Patient metadata and retained multimodal data.** Patient-level metadata for the glioblastoma cohort included in the study. One invasive-region section was excluded after quality control, resulting in 19 retained tissue sections.

| ID | Age | MGMT | IDH | p53 | GFAP | KI67 | ATRX | Overall Survival | Survival event | Region availability | Cells Tumor | Cells Invasive | Cells total |
|---|---|---|---|---|---|---|---|---|---|---|---|---|---|
| 1 | 60 | + | WT | - | + | + | + | 40.77 | alive | Tumor and Invasive | 8244 | 13374 | 21618 |
| 2 | 32 | - | WT | + | + | + | - | 28.27 | alive | Tumor and Invasive | 16530 | 8948 | 25478 |
| 3 | 52 | - | WT | + | + | + | + | 13 | death | Tumor and Invasive | 9371 | 6007 | 15378 |
| 4 | 69 | + | WT | + | + | + | + | 22.41 | death | Tumor and Invasive | 9170 | 4329 | 13499 |
| 5 | 65 | + | WT | + | + | + | + | 26.31 | alive | Tumor and Invasive | 1428 | 13623 | 15051 |
| 6 | 64 | + | WT | + | + | + | + | 34.16 | death | Tumor and Invasive | 9447 | 8250 | 17697 |
| 7 | 67 | - | WT | - | + | + | - | 14.06 | death | Tumor and Invasive | 18327 | 1817 | 20144 |
| 8 | 65 | - | WT | + | + | + | + | 15.3 | death | Tumor only | 18666 | 0 | 18666 |
| 9 | 58 | - | WT | + | + | + | + | 9.26 | death | Tumor and Invasive | 17391 | 13906 | 31297 |
| 10 | 73 | + | WT | + | + | + | + | 30.1 | alive | Tumor and Invasive | 14133 | 5176 | 19309 |

**Supplementary Table S3. Patient-level cell counts by anatomical region and cell type.** Table showing the number of annotated cells per patient, anatomical region, and cell type after Xenium processing and quality filtering. For each retained tumor-core and invasive-region section, counts are reported for astrocytes, endothelial cells, excitatory neurons, GSCs, inhibitory neurons, MES-like cells, OPCs, oligodendrocytes, TAMs, and T cells, together with the total number of cells analyzed per section.

| ID | zone | Astrocytes | Endothelial | Excitatory Neu. | GSC | Inhibitory Neu. | MES | OPC | Oligodendr. | TAM | T Cell | Total | Area (mm$^2$) |
|---|---|---|---|---|---|---|---|---|---|---|---|---|---|
| 1 | Tumor | 4177 | 404 | 14 | 621 | 26 | 443 | 392 | 905 | 510 | 752 | 8244 | 4.91 |
| 1 | Invasive | 4588 | 683 | 343 | 1303 | 120 | 403 | 1204 | 3333 | 536 | 861 | 13374 | 5.27 |
| 2 | Tumor | 1117 | 641 | 16 | 558 | 47 | 78 | 10267 | 2544 | 200 | 1062 | 16530 | 6.28 |
| 2 | Invasive | 1168 | 261 | 14 | 234 | 26 | 43 | 3714 | 2881 | 161 | 446 | 8948 | 3.84 |
| 3 | Tumor | 3954 | 1104 | 277 | 833 | 288 | 403 | 651 | 180 | 702 | 979 | 9371 | 1.66 |
| 3 | Invasive | 3555 | 314 | 92 | 483 | 103 | 156 | 579 | 386 | 171 | 168 | 6007 | 1.55 |
| 4 | Tumor | 1331 | 1341 | 2 | 778 | 10 | 974 | 411 | 141 | 714 | 3468 | 9170 | 1.99 |
| 4 | Invasive | 1256 | 193 | 6 | 208 | 11 | 52 | 342 | 1609 | 204 | 448 | 4329 | 2.26 |
| 5 | Tumor | 1121 | 33 | 24 | 32 | 16 | 38 | 50 | 27 | 73 | 14 | 1428 | 1.46 |
| 5 | Invasive | 4053 | 682 | 30 | 785 | 84 | 187 | 1001 | 5283 | 508 | 1010 | 13623 | 7.39 |
| 6 | Tumor | 1243 | 727 | 97 | 871 | 90 | 1399 | 184 | 95 | 1126 | 3615 | 9447 | 3.15 |
| 6 | Invasive | 2223 | 413 | 230 | 428 | 130 | 219 | 506 | 2446 | 580 | 1075 | 8250 | 5.14 |
| 7 | Tumor | 8828 | 1442 | 943 | 1860 | 796 | 684 | 1192 | 607 | 890 | 1085 | 18327 | 6.97 |
| 7 | Invasive | 460 | 136 | 142 | 62 | 48 | 33 | 187 | 545 | 62 | 142 | 1817 | 1.48 |
| 8 | Tumor | 5765 | 1233 | 537 | 3547 | 201 | 3356 | 2227 | 231 | 416 | 1153 | 18666 | 4.84 |
| 9 | Tumor | 13817 | 376 | 94 | 1816 | 129 | 202 | 483 | 162 | 189 | 123 | 17391 | 2.65 |
| 9 | Invasive | 8869 | 748 | 335 | 1552 | 271 | 275 | 728 | 583 | 294 | 251 | 13906 | 4.47 |
| 10 | Tumor | 4323 | 1067 | 66 | 1088 | 50 | 483 | 935 | 1280 | 1322 | 3519 | 14133 | 4.95 |
| 10 | Invasive | 1768 | 248 | 8 | 263 | 11 | 112 | 457 | 1530 | 217 | 562 | 5176 | 2.26 |

**Supplementary Table S4. Marker genes used for gene program annotation.** The table lists the marker genes used to support annotation of the following gene programs: proliferation, hypoxia, ECM remodeling, TAMs phagocytosis, T cell infiltration and lipid synthesis.

| Gene Program | Marker list |
|---|---|
| Proliferation | CCNA1, CCNB2, CDK1, CENPF, KIT, MKI67, PCNA, TOP2A |
| Hypoxia | ANGPT1, CXCR4, IGFBP3, LOX, MGST1, RNASET2 |
| ECM Remodeling | ADAMTS12, ADAMTS16, ADAMTS3, BCAN, CEMIP, CHODL, COL12A1, DCN, LAMA2, VCAN |
| TAMs phagocytosis | AIF1, CAPG, CD14, CD163, CD68, CTSS, CX3CR1, CYTIP, FCER1G, FCGR3A,GPR183, GPR34, ITGB2, LY86, MS4A6A, RGS10 |
| T Cell infiltration | CD2, CD3G, CD4, CD48, CD52, GNLY, IL7R, KLRB1, NKG7, STXBP2, TRAC |
| Lipid Synthesis | APOE, B4FALNT1, CAV1, CD36, ELOVL2, HILPDA, HS3ST2, HS3ST4, IDH1, IDH2, IGFBP5, NOTCH1, NR2F2, PLD5, SLC26A4, SPHKAP, STAT3, SULF1, TGFB1, TGFB2, UGT8 |

**Supplementary Table S5. Annotated lipid features used for MALDI-MSI lipidomic analyses.** Lipid annotations used for class-level, species-level, spatial, and lipid–gene coupling analyses. For each feature, the table reports the LC-MS/MS-derived m/z, lipid class, lipid abbreviation, chain-level annotation from Lipostar/LIPID MAPS database matching, detected adduct, MALDI-MSI feature m/z, matched MALDI m/z, and matching mass error in parts per million (ppm). These annotations were used to assign MALDI-MSI features to lipid classes and individual lipid species.

| LC-MS/MS_mz | lipid_class | abbreviation | abbreviation_chains (Lipostar LIPID MAPS DB) | adduct | feature | maldi_mz | ppm_match |
|---|---|---|---|---|---|---|---|
| 372.312 | CAR | CAR | 14:0 | [M+H]+ | 372.311 | 372.311 | -1.9374 |
| 398.327 | CAR | CAR | 16:1 | [M+H]+ | 398.329 | 398.329 | 3.9049 |
| 424.3425 | CAR | CAR | 18:2 | [M+H]+ | 424.34 | 424.34 | -5.9089 |
| 426.3581 | CAR | CAR | 18:1 | [M+H]+ | 426.36 | 426.36 | 4.388 |
| 428.374 | CAR | CAR | 18:0 | [M+H]+ | 428.369 | 428.369 | -12.6 |
| 454.2932 | O-LPE | LPE O-16:1 | LPE O-16:1;O | [M+H]+ | 454.289 | 454.289 | -10.1872 |
| 468.3091 | LPC | LPC 14:0 | LPC 14:0 | [M+H]+ | 468.311 | 468.311 | 4.9157 |
| 494.3247 | LPC | LPC 16:1 | LPC 16:1 | [M+H]+ | 494.32 | 494.32 | -9.6381 |
| 496.3402 | LPC | LPC 16:0 | LPC 16:0 | [M+H]+ | 496.34 | 496.34 | -0.3986 |
| 502.2933 | LPE | LPE 20:4 | LPE 20:4 | [M+H]+ | 502.3 | 502.3 | 13.3381 |
| 518.3224 | LPC | LPC 16:0 | LPC 16:0 | [M+Na]+ | 518.32 | 518.32 | -4.5167 |
| 520.3404 | LPC | LPC 18:2 | LPC 18:2 | [M+H]+ | 520.34 | 520.34 | -0.7654 |
| 522.3561 | LPC | LPC 18:1 | LPC 18:1 | [M+H]+ | 522.36 | 522.36 | 7.481 |
| 524.3718 | LPC | LPC 18:0 | LPC 18:0 | [M+H]+ | 524.369 | 524.369 | -6.1582 |
| 526.2936 | LPE | LPE 22:6 | LPE 22:6 | [M+H]+ | 526.3 | 526.3 | 12.1633 |
| 538.5195 | Cer | Cer 34:1 | Cer d18:1_16:0 | [M+H]+ | 538.511 | 538.511 | -14.9906 |
| 544.3405 | LPC | LPC 20:4 | LPC 20:4 | [M+H]+ | 544.34 | 544.34 | -0.923 |
| 546.3558 | LPC | LPC 20:3 | LPC 20:3 | [M+H]+ | 546.36 | 546.36 | 7.6873 |
| 564.5352 | Cer | Cer 36:2 | Cer d18:2_18:0 | [M+H]+ | 564.529 | 564.529 | -11.7416 |
| 566.5508 | Cer | Cer 36:1 | Cer d18:1_18:0 | [M+H]+ | 566.549 | 566.549 | -3.9336 |
| 594.5824 | Cer | Cer 38:1 | Cer d18:1_20:0 | [M+H]+ | 594.58 | 594.58 | -4.0496 |
| 622.6135 | Cer | Cer 40:1 | Cer d16:1_24:0 | [M+H]+ | 622.609 | 622.609 | -7.9369 |
| 634.6137 | Cer | Cer 41:2 | Cer d18:2_23:0 | [M+H]+ | 634.62 | 634.62 | 9.9273 |
| 636.6294 | Cer | Cer 41:1 | Cer d18:1_23:0 | [M+H]+ | 636.631 | 636.631 | 3.1845 |
| 642.6032 | DG | DG 36:0 | DG 18:0_18:0 | [M+NH4]+ | 642.609 | 642.609 | 8.3586 |
| 648.629 | Cer | Cer 42:2 | Cer d18:1_24:1 | [M+H]+ | 648.631 | 648.631 | 3.7613 |
| 650.6447 | Cer | Cer 42:1 | Cer d18:1_24:0 | [M+H]+ | 650.64 | 650.64 | -7.1128 |
| 662.6452 | Cer | Cer 43:2 | Cer d18:2_25:0 | [M+H]+ | 662.64 | 662.64 | -7.8473 |
| 671.5736 | CE | CE 18:2 | CE 18:2 | [M+Na]+ | 671.573 | 671.573 | -1.0114 |
| 674.5116 | P-PE | PE P-32:1 | PE P-16:0_16:1 | [M+H]+ | 674.521 | 674.521 | 13.9973 |
| 676.5274 | P-PE | PE P-32:0 | PE P-16:0_16:0 | [M+H]+ | 676.529 | 676.529 | 1.6686 |
| 684.6138 | TG | TG 38:0 | TG 12:0_12:0_14:0 | [M+NH4]+ | 684.608 | 684.608 | -8.6694 |
| 686.5718 | DG | DG 40:6 | DG 18:0_22:6 | [M+NH4]+ | 686.571 | 686.571 | -0.541 |
| 688.6026 | CE | CE 20:5 | CE 20:5 | [M+NH4]+ | 688.609 | 688.609 | 8.6721 |
| 690.543 | O-PC | PC O-30:1 | PC O-14:0_16:1 | [M+H]+ | 690.548 | 690.548 | 7.8415 |
| 695.5737 | CE | CE 20:4 | CE 20:4 | [M+Na]+ | 695.571 | 695.571 | -4.1909 |
| 697.5259 | SM | SM 32:1 | SM d16:1_16:0 | [M+Na]+ | 697.522 | 697.522 | -6.2511 |
| 697.5891 | CE | CE 20:3 | CE 20:3 | [M+Na]+ | 697.58 | 697.58 | -13.3006 |
| 702.5428 | P-PE | PE P-34:1 | PE P-16:0_18:1 | [M+H]+ | 702.549 | 702.549 | 8.2152 |
| 704.523 | PC | PC 30:1 | PC 12:0_18:1 | [M+H]+ | 704.52 | 704.52 | -4.3699 |
| 705.5904 | SM | SM 34:0 | SM d16:0_18:0 | [M+H]+ | 705.58 | 705.58 | -14.8184 |
| 706.538 | PC | PC 30:0 | PC 10:0_20:0 | [M+H]+ | 706.54 | 706.54 | 2.6965 |
| 716.5223 | PE | PE 34:2 | PE 12:0_22:2 | [M+H]+ | 716.529 | 716.529 | 8.7682 |
| 717.5907 | SM | SM 35:1 | SM d18:1_17:0 | [M+H]+ | 717.58 | 717.58 | -14.8541 |
| 719.5697 | SM | SM 34:1 | SM d18:1_16:0(2OH) | [M+H]+ | 719.56 | 719.56 | -13.4802 |
| 720.5541 | PE | PE 34:0 | PE 12:0_22:0 | [M+H]+ | 720.56 | 720.56 | 8.1882 |
| 723.5417 | SM | SM 34:2 | SM d16:1_18:1 | [M+Na]+ | 723.541 | 723.541 | -1.5339 |
| 725.5565 | SM | SM 34:1 | SM d16:1_18:0 | [M+Na]+ | 725.549 | 725.549 | -10.7299 |
| 726.5435 | O-PE | PE O-36:4 | PE O-16:0_20:4 | [M+H]+ | 726.549 | 726.549 | 7.1754 |
| 728.5198 | PC | PC 30:0 | PC 13:0_17:0 | [M+Na]+ | 728.521 | 728.521 | 1.2246 |
| 729.5908 | SM | SM 36:2 | SM d16:1_20:1 | [M+H]+ | 729.591 | 729.591 | 0.938 |
| 730.5382 | PC | PC 32:2 | PC 12:0_20:2 | [M+H]+ | 730.529 | 730.529 | -13.0596 |
| 734.5694 | PC | PC 32:0 | PC 10:0_22:0 | [M+H]+ | 734.571 | 734.571 | 2.7459 |
| 738.5075 | PE | PE 36:5 | PE 16:0_20:5 | [M+H]+ | 738.511 | 738.511 | 5.2727 |
| 739.5726 | SM | SM 35:1 | SM d18:1_17:0 | [M+Na]+ | 739.571 | 739.571 | -1.5852 |
| 740.5223 | PE | PE 36:4 | PE 16:0_20:4 | [M+H]+ | 740.522 | 740.522 | -0.658 |
| 742.5377 | PE | PE 36:3 | PE 14:1_22:2 | [M+H]+ | 742.543 | 742.543 | 6.9522 |
| 744.5537 | PE | PE 36:2 | PE 18:1_18:1 | [M+H]+ | 744.56 | 744.56 | 8.4669 |
| 746.5694 | PE | PE 36:1 | PE 18:0_18:1 | [M+H]+ | 746.577 | 746.577 | 10.5936 |
| 748.5848 | PC | PC 33:0 | PC 16:0_17:0 | [M+H]+ | 748.58 | 748.58 | -6.4062 |
| 752.5585 | P-PE | PE P-38:4 | PE P-18:0_20:4 | [M+H]+ | 752.56 | 752.56 | 1.7559 |
| 753.5876 | SM | SM 36:1 | SM d16:1_20:0 | [M+Na]+ | 753.58 | 753.58 | -10.0931 |
| 754.5388 | PC | PC 34:4 | PC 12:0_22:4 | [M+H]+ | 754.54 | 754.54 | 1.5164 |
| 756.5508 | PC | PC 32:0 | PC 10:0_22:0 | [M+Na]+ | 756.54 | 756.54 | -14.2637 |
| 756.5905 | O-PE | PE O-38:3 | PE O-18:0_20:3 | [M+H]+ | 756.598 | 756.598 | 10.3083 |
| 758.569 | PC | PC 34:2 | PC 12:0_22:2 | [M+H]+ | 758.571 | 758.571 | 3.1844 |
| 759.6374 | SM | SM 38:1 | SM d16:1_22:0 | [M+H]+ | 759.631 | 759.631 | -8.5718 |
| 760.5846 | PC | PC 34:1 | PC 12:0_22:1 | [M+H]+ | 760.591 | 760.591 | 8.9726 |
| 761.6534 | SM | SM 38:0 | SM d16:0_22:0 | [M+H]+ | 761.643 | 761.643 | -13.8011 |
| 762.5075 | PE | PE 38:7 | PE 16:1_22:6 | [M+H]+ | 762.511 | 762.511 | 5.0644 |
| 762.5961 | PC | PC 34:0 | PC 10:0_24:0 | [M+H]+ | 762.591 | 762.591 | -6.1408 |
| 766.5384 | PE | PE 38:5 | PE 18:0_20:5 | [M+H]+ | 766.54 | 766.54 | 2.0733 |
| 768.5536 | PE | PE 38:4 | PE 18:0_20:4 | [M+H]+ | 768.549 | 768.549 | -6.5569 |
| 770.5696 | PC | PC 35:3 | PC 15:0_20:3 | [M+H]+ | 770.573 | 770.573 | 4.6265 |
| 772.5845 | PC | PC 35:2 | PC 13:0_22:2 | [M+H]+ | 772.58 | 772.58 | -5.9127 |
| 774.6004 | PC | PC 35:1 | PC 13:0_22:1 | [M+H]+ | 774.6 | 774.6 | -0.5183 |
| 775.5933 | PC | PC 33:1 | PC 15:0_18:1 | [M+Na]+ | 775.6 | 775.6 | 8.6364 |
| 778.5386 | PC | PC 36:6 | PC 14:0_22:6 | [M+H]+ | 778.54 | 778.54 | 1.7036 |
| 780.5509 | PC | PC 34:2 | PC 12:0_22:2 | [M+Na]+ | 780.549 | 780.549 | -3.0664 |
| 780.5902 | P-PE | PE P-40:4 | PE P-18:0_22:4 | [M+H]+ | 780.596 | 780.596 | 7.7294 |
| 781.6193 | SM | SM 38:1 | SM d16:1_22:0 | [M+Na]+ | 781.608 | 781.608 | -14.1413 |
| 782.5694 | PC | PC 36:4 | PC 14:0_22:4 | [M+H]+ | 782.571 | 782.571 | 2.5841 |
| 783.6366 | SM | SM 40:3 | SM d18:2_22:1 | [M+H]+ | 783.631 | 783.631 | -6.8802 |
| 784.5829 | PC | PC 34:0 | PC 10:0_24:0 | [M+Na]+ | 784.571 | 784.571 | -14.6254 |
| 786.6001 | PC | PC 36:2 | PC 14:0_22:2 | [M+H]+ | 786.591 | 786.591 | -11.0167 |
| 787.6688 | SM | SM 40:1 | SM d16:1_24:0 | [M+H]+ | 787.66 | 787.66 | -11.3332 |
| 788.6161 | PC | PC 36:1 | PC 14:0_22:1 | [M+H]+ | 788.609 | 788.609 | -9.5391 |
| 790.5375 | PE | PE 40:7 | PE 18:1_22:6 | [M+H]+ | 790.54 | 790.54 | 3.1568 |
| 790.6288 | PC | PC 36:0 | PC 11:0_25:0 | [M+H]+ | 790.62 | 790.62 | -11.1128 |
| 792.5536 | PE | PE 40:6 | PE 18:0_22:6 | [M+H]+ | 792.56 | 792.56 | 8.057 |
| 796.5849 | PC | PC 37:4 | PC 15:0_22:4 | [M+H]+ | 796.58 | 796.58 | -6.2784 |

| LC-MS/MS_mz | lipid_class | abbreviation | abbreviation_chains (Lipostar LIPID MAPS DB) | adduct | feature | maldi_mz | ppm_match |
|---|---|---|---|---|---|---|---|
| 796.666 | GalCer | GalCer 41:2 | GalCer d18:2_23:0 | [M+H]+ | 796.671 | 796.671 | 6.8064 |
| 798.5993 | PE | PE 40:3 | PE 18:1_22:2 | [M+H]+ | 798.59 | 798.59 | -11.315 |
| 799.6686 | SM | SM 41:2 | SM d17:1_24:1 | [M+H]+ | 799.66 | 799.66 | -10.786 |
| 800.6611 | GlcCer | GlcCer 40:1 | GlcCer d18:1_22:0(2OH[R]) | [M+H]+ | 800.66 | 800.66 | -1.5817 |
| 801.6846 | SM | SM 41:1 | SM d16:1_25:0 | [M+H]+ | 801.68 | 801.68 | -5.7331 |
| 802.536 | PC | PC 36:5 | PC 14:0_22:5 | [M+Na]+ | 802.54 | 802.54 | 4.9429 |
| 802.6321 | PC | PC 37:1 | PC 15:0_22:1 | [M+H]+ | 802.62 | 802.62 | -14.9437 |
| 804.5509 | PC | PC 36:4 | PC 14:0_22:4 | [M+Na]+ | 804.54 | 804.54 | -13.5341 |
| 806.5697 | PC | PC 38:6 | PC 16:0_22:6 | [M+H]+ | 806.571 | 806.571 | 2.1486 |
| 807.6349 | SM | SM 40:2 | SM d16:1_24:1 | [M+Na]+ | 807.63 | 807.63 | -5.7403 |
| 808.582 | PC | PC 36:2 | PC 14:0_22:2 | [M+Na]+ | 808.571 | 808.571 | -13.0757 |
| 809.651 | SM | SM 40:1 | SM d18:1_22:0 | [M+Na]+ | 809.648 | 809.648 | -3.1077 |
| 810.6001 | PC | PC 38:4 | PC 16:0_22:4 | [M+H]+ | 810.6 | 810.6 | -0.1069 |
| 811.6681 | SM | SM 42:3 | SM d18:2_24:1 | [M+H]+ | 811.671 | 811.671 | 4.1449 |
| 813.6843 | SM | SM 42:2 | SM d18:1_24:1 | [M+H]+ | 813.68 | 813.68 | -5.2859 |
| 814.5377 | PE | PE 42:9 | PE 20:3_22:6 | [M+H]+ | 814.54 | 814.54 | 2.8188 |
| 814.6771 | GlcCer | GlcCer 39:1 | GlcCer d16:1(15Me)_23:0(22Me)(2OH) | [M+H]+ | 814.68 | 814.68 | 3.5006 |
| 815.7001 | SM | SM 42:1 | SM d18:0_24:1 | [M+H]+ | 815.691 | 815.691 | -10.6714 |
| 820.5842 | PC | PC 39:6 | PC 17:0_22:6 | [M+H]+ | 820.58 | 820.58 | -5.4032 |
| 821.6504 | SM | SM 41:2 | SM d17:1_24:1 | [M+Na]+ | 821.648 | 821.648 | -2.5948 |
| 822.637 | O-PC | PC O-40:5 | PC O-18:0_22:5 | [M+H]+ | 822.64 | 822.64 | 3.8054 |
| 823.6665 | SM | SM 41:1 | SM d16:1_25:0 | [M+Na]+ | 823.66 | 823.66 | -7.9246 |
| 828.5511 | PC | PC 38:6 | PC 16:0_22:6 | [M+Na]+ | 828.549 | 828.549 | -2.9428 |
| 828.6917 | GlcCer | GlcCer 42:1 | GlcCer d18:1_24:0(2OH[R]) | [M+H]+ | 828.7 | 828.7 | 10.0255 |
| 829.7159 | SM | SM 43:1 | SM d18:1_25:0 | [M+H]+ | 829.711 | 829.711 | -5.3891 |
| 832.5821 | PC | PC 38:4 | PC 16:0_22:4 | [M+Na]+ | 832.571 | 832.571 | -12.8314 |
| 833.6517 | SM | SM 42:3 | SM d18:2_24:1 | [M+Na]+ | 833.649 | 833.649 | -3.7948 |
| 834.598 | PC | PC 38:3 | PC 16:1_22:2 | [M+Na]+ | 834.591 | 834.591 | -7.8626 |
| 835.6662 | SM | SM 42:2 | SM d18:1_24:1 | [M+Na]+ | 835.66 | 835.66 | -7.431 |
| 836.523 | PE | PE 44:12 | PE 22:6_22:6 | [M+H]+ | 836.529 | 836.529 | 6.6627 |
| 837.6805 | SM | SM 42:1 | SM d18:0_24:1 | [M+Na]+ | 837.671 | 837.671 | -10.8741 |
| 841.7155 | SM | SM 44:2 | SM d18:1_26:1 | [M+H]+ | 841.711 | 841.711 | -4.8381 |
| 842.7233 | TG | TG 50:5 | TG 14:0_18:2_18:3 | [M+NH4]+ | 842.72 | 842.72 | -3.9172 |
| 843.7306 | SM | SM 44:1 | SM d18:0_26:1 | [M+H]+ | 843.72 | 843.72 | -12.7213 |
| 844.7389 | TG | TG 50:4 | TG 14:0_16:0_20:4 | [M+NH4]+ | 844.729 | 844.729 | -12.2738 |
| 848.654 | O-PC | PC O-42:6 | PC O-20:0_22:6 | [M+H]+ | 848.649 | 848.649 | -6.3966 |
| 849.6822 | SM | SM 43:2 | SM d17:1_26:1 | [M+Na]+ | 849.671 | 849.671 | -12.677 |
| 850.6713 | P-PC | PC P-42:4 | PC P-20:0_22:4 | [M+H]+ | 850.671 | 850.671 | 0.1511 |
| 854.5668 | PC | PC 40:7 | PC 18:1_22:6 | [M+Na]+ | 854.56 | 854.56 | -7.8295 |
| 856.5821 | PC | PC 40:6 | PC 18:0_22:6 | [M+Na]+ | 856.571 | 856.571 | -12.459 |
| 858.5978 | PC | PC 40:5 | PC 18:0_22:5 | [M+Na]+ | 858.591 | 858.591 | -7.4271 |
| 858.7546 | TG | TG 51:4 | TG 15:1_18:0_18:3 | [M+NH4]+ | 858.751 | 858.751 | -3.6882 |
| 870.6946 | PC | PC 42:2 | PC 16:0_26:2 | [M+H]+ | 870.691 | 870.691 | -3.6022 |
| 872.7104 | PC | PC 42:1 | PC 18:0_24:1 | [M+H]+ | 872.7 | 872.7 | -11.9407 |
| 881.5153 | PI | PI 36:4 | PI 14:0_22:4 | [M+Na]+ | 881.511 | 881.511 | -4.3757 |
| 881.7565 | TG | TG 52:2 | TG 16:0_18:1_18:1 | [M+Na]+ | 881.751 | 881.751 | -5.75 |
| 885.5455 | PI | PI 36:2 | PI 14:0_22:2 | [M+Na]+ | 885.549 | 885.549 | 3.5171 |
| 887.5642 | PI | PI 38:4 | PI 18:0_20:4 | [M+H]+ | 887.571 | 887.571 | 8.0744 |
| 896.7102 | PC | PC 44:3 | PC 22:1_22:2 | [M+H]+ | 896.7 | 896.7 | -11.4131 |
| 898.7261 | PC | PC 44:2 | PC 22:0_22:2 | [M+H]+ | 898.72 | 898.72 | -7.0332 |
| 903.7404 | TG | TG 54:5 | TG 18:1_18:1_18:3 | [M+Na]+ | 903.729 | 903.729 | -13.0972 |
| 907.772 | TG | TG 54:3 | TG 18:1_18:1_18:1 | [M+Na]+ | 907.76 | 907.76 | -13.2176 |
| 909.5459 | PI | PI 38:4 | PI 16:0_22:4 | [M+Na]+ | 909.54 | 909.54 | -6.5187 |
| 920.7078 | PC | PC 44:2 | PC 22:0_22:2 | [M+Na]+ | 920.7 | 920.7 | -8.4718 |
| 935.803 | TG | TG 56:3 | TG 16:0_18:3_22:0 | [M+Na]+ | 935.791 | 935.791 | -12.3603 |
| 948.7393 | PC | PC 46:2 | PC 23:1_23:1 | [M+Na]+ | 948.729 | 948.729 | -11.3082 |
